# Assessing tissue-specific gene expression of essential genes from human and mouse

**DOI:** 10.1101/2023.12.21.572731

**Authors:** Huiwen Zheng, Atefeh Taherian Fard, Jessica C Mar

## Abstract

A gene satisfies the definition of essentiality when the loss of its function compromises an organism’s viability. Identifying essential genes is useful for understanding the core components that regulate a biological system and ensure its survival. Advances in gene editing techniques like CRISPR-Cas9 generate the capacity to comprehensively interrogate a genome to elucidate what genes are essential. However, these techniques are often applied in the context of a single cell line and even when studies have collated essential gene sets across multiple cell lines, this information is rarely probed at a level that incorporates multiple cell and tissue types. The recent availability of large-scale single-cell RNA-sequencing (scRNA-seq) atlases provides an unprecedented opportunity to investigate the distribution of essential gene expression in cell and tissue types.

Here, we leverage information contained in benchmarking datasets, single cell tissue atlases, and databases of essential genes, and develop a computational method, scEssentials, which uses a statistical framework to report on the robustness and specificity of essential genes in human and mouse across multiple cell types. Using scEssentials, both mouse and human models showed consistently high in expression and exhibit limited variably across more than 60 cell types. We also demonstrate a substantial number of significantly correlated gene pairs within scEssentials, which produce densely connected co-expression networks with functional annotation. Furthermore, we showed high frequencies of scEssentials across 200 pathways. Finally, we develop a score to quantify the relative essentiality of genes within scEssentials, which further validates with significant association with gene mutation frequency and chromatin accessibility.

Using the heterogeneous ageing process, we demonstrate the application of scEssentials and their robust gene expression profile. Only one-fifth of scEssentials showed significant ageing-related differential expression among three age groups, occurring primarily in muscle satellite cells of varying tissue origins and highly interacting brain cells. Collectively, the robustness of scEssentials serves as a reference for analysing scRNA-seq data and provides insight into the heterogeneous nature of biological processes such as ageing.

## Introduction

Gene essentiality is a valuable concept for delineating the rules of a biological system because removing essential genes exposes the core genetic machinery that an organism requires to survive (2). While essential genes are useful for understanding genetics, it is consideration of their gene expression of that is valuable for functional genomics. In this study, we develop a computational approach to investigate the cell type specific expression of essential genes in a robust and reliable way for human and mouse data. We leverage the vast amount of information pertaining to essential genes that currently exist and combine this with single cell tissue atlases to provide sets of essential genes that are characterised by their expression level, gene expression variability, co-expression and other properties.

Typically, essential genes are identified without consideration to the specificity or context of cell or tissue types in which they are expressed in. The experimental methods that are used to identify essential genes are rarely conducted at single-cell resolution and in tissue-wide manner. Advancements in molecular techniques have contributed to the availability of comprehensive databases of essential genes that have been validated for different organisms. Recently, Luo *et al*. updated the DEG15 database that summarised an increased number of eukaryotic and prokaryotic essential genes identified through transposon-insertion sequencing, CRISPR/Cas9 and single-gene knockout experiments (2). The most significant progress has been made in human essential genes, primarily due to the research studies to investigate the functions of essential gene function in different biological systems (3, 4). Even when comparing human and mouse, the set of homologous essential genes between these species had minimum overlap which points to change in essentiality during evolution (5). Additionally, genes identified as essential in human cell lines or knockout mice may be distinct from those in living humans (6). All the evidence improves the knowledge of the essentiality conservation happens at the functional level, rather than the gene level (5, 7).

To qualify as an essential gene, these genes must satisfy experimental validations of their essentiality such as through CRISPR /Cas9 or gene-knockout experiments. In systems biology, it is well known that some essential genes often overlap with housekeeping genes, ribosomes, and stably expressed genes because all of these gene sets function in roles that are critical to the survival of the organism. This overlap between gene sets results from the redundancy that is built into how biological systems are wired. For example, housekeeping genes are regarded as essential for cellular maintenance but not always be stably expressed or essential across multiple species (8). Examples exist where some ribosomes are essential genes for all organisms but their expression patterns can be altered in tumour cells (9). Unlike ribosomes and essential genes which are based on defined based on functional properties, stably expressed genes (SEGs) are commonly identified using statistical modelling based on their pattern of gene expression. SEGs are often used for normalisation or the standardisation of signals, and some of these genes also play an essential role in cellular survival independent of the experimental conditions. The stability properties that lead to SEGs are not usually validated experimentally and hence a gene may function as a SEG only in the context of the dataset that was used to identify these genes (10). While examples exist where essential genes are also known to serve as a housekeeping gene that also has stable gene expression, it is important to recognize that these three definitions are unique. For instance, studies in humans demonstrated that housekeeping genes exhibit greater evolutionary conservation than essential genes do (11) while essential genes are associated with greater lethality and not associated with human diseases (12). Moreover, the substantial difference between human orthologs of mouse essential genes and human essential genes emphasis the need of generating organism-specific essential genes. Therefore, careful considerations are required before using any of the gene sets as the reference to investigate the consequences of such gene expression changes in different organisms.

Given that the identify of many essential genes have been established, more recently, efforts have been made to extend this information by investigating their functional properties with respect to regulating an overall system. Li *et al*. summarised the topological characteristics of essential genes in protein-protein interaction networks and their related biological information (13). Previous studies have investigated the characteristics of the essential genes at the transcriptome level and also revealed the important functional roles in the epigenetic landscape of normal cells (5). These studies provide a foundation for identifying essential genes that are important to maintain cell viability. Understanding how essential genes are expressed in the context of a single cell type is valuable because cell type identity is driven by gene expression programs. Given that only a fraction of genes is expressed in a cell type, investigating whether essential genes are robust in their expression profiles and expressed across multiple cell types can extend our knowledge of how essential genes are expressed within the context of an entire organism and provide computational inferences of which essential genes are more critical to cell survival than others (?). Advances in sequencing technologies have made capturing transcriptomes at single-cell resolution a reality. As such, we leveraged the available single cell atlases to investigate how essential genes were expressed within cell type in human and mouse organisms. With the low capture efficiency and sequencing limitation for scRNA-seq data, it is not unexpected to observe missing genes in the sequencing data. The presence of these missing genes may be a direct result of the limitation of scRNA-seq technology and platform specific effects should also be taken into account in the analysis. Additionally, the scope of assay must also be accommodated since data collected via *in vivo* assays is susceptible to higher amounts of noise than *in vitro* data. This experimental context should be acknowledged when considering what studies were used to identify essential genes.

Therefore, investigating the characteristics of essential genes at the cell-type level with large-scale atlases from scRNA-seq data like Tabula-Muris (TM) and Tabula-Sapiens (TS) is crucial (14, 15). To the best of our knowledge, it is the first evaluation of essential genes at the single-cell level in human and mouse. Our study aims to generate gene sets that satisfy computational properties of essential genes across cell types in normal conditions. These gene sets represent utility for future perturbation studies where the loss of gene expression for these specific genes are more likely to lead to severe consequences.

In this study, we first measure which genes are likely to be essential based on properties with respect to robustness, corresponding biological functions and co-expression connectivity. We use over 100 cell types and 10 different sequencing platforms to identify a set of essential genes that satisfy quality control and reliability detection for 1single-cell RNA-seq data using a computational approach termed scEssentials.. These pre-ranked genes can be applied in any scRNA-seq dataset to either normalise the data or identify the gene expression changes. Next, we evaluated the characteristics of the scEssentials and their robustness in multi-omics data. High expression levels and robust essentiality scores were found highly associated with other molecular levels. Lastly, we demonstrated the stable expression for the scEssentials genes during ageing to show the characteristics of scEssentials in the dynamic and heterogeneous biological processes.

## Methods

### Single-cell RNA-sequencing (scRNA-seq) datasets used for evaluating expression of essential genes

Single cell RNA-seq datasets were sources from human and mouse to evaluate the specificity and reliability of essential gene expression. For specificity, we used the Tabula Muris (TM) and Tabula Sapiens (TS) which are single cell atlases that cover a broad range of tissue and cell types from mouse and human, respectively. [Details of data availability, download, version, etc.] Our study used the data generated by FACs SmartSeq2 technology since these datasets are less sparsity than the Droplet-based options in TM (and TS?). We used data from the young groups (3 months for TM and <40 years for TS) with 7 mice and 5 donors, respectively. Only cell types that have more than 100 cells were included which resulted in 68 unique tissue-cell-types for TM and 53 unique tissue-cell-types for TS.

Two benchmarking datasets were used to evaluate reliability of expression by assessing technical variation. The mouse embryonic stem cells (mESCs) that have been sequenced by CELseq2, Dropseq, MARSeq, SCRBseq, Smartseq and Smartseq2 (16) and a matched control ‘normal’ B lymphocyte line from breast cancer patients that have been sequenced by 10x Genomics Chromium, Fluidigm C1, Fluidigm C1 HT and Takara Bio ICELL8 (17) as listed in Supplementary Table 1.

### Source of essential gene information

Lists of essential genes identified for human and mouse were downloaded from the DEG15 database (2). A total of 2354 mouse essential genes and 8988 human essential genes were curated through different experimental techniques, mainly via CRISPR/Cas9 and single-gene knockout validations.

### Developing a computational method, scEssentials, **to ass**ess essential gene expression

Because single cell RNA-sequencing experiments typically have higher dropout rates and technical limitations that affect scale, some of these essential genes were not detected in scRNA-seq data. Therefore, we imposed a series of selection criteria to ensure the scEssentials generalisability.

First, we used two benchmarking datasets to ensure the detectability of scEssentials across major sequencing platforms, so that for any given essential genes *x*_*m*_ at least 3 non-zero reads in every sequencing method (Figure 1), denoted as:

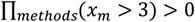

**Figure 1.**
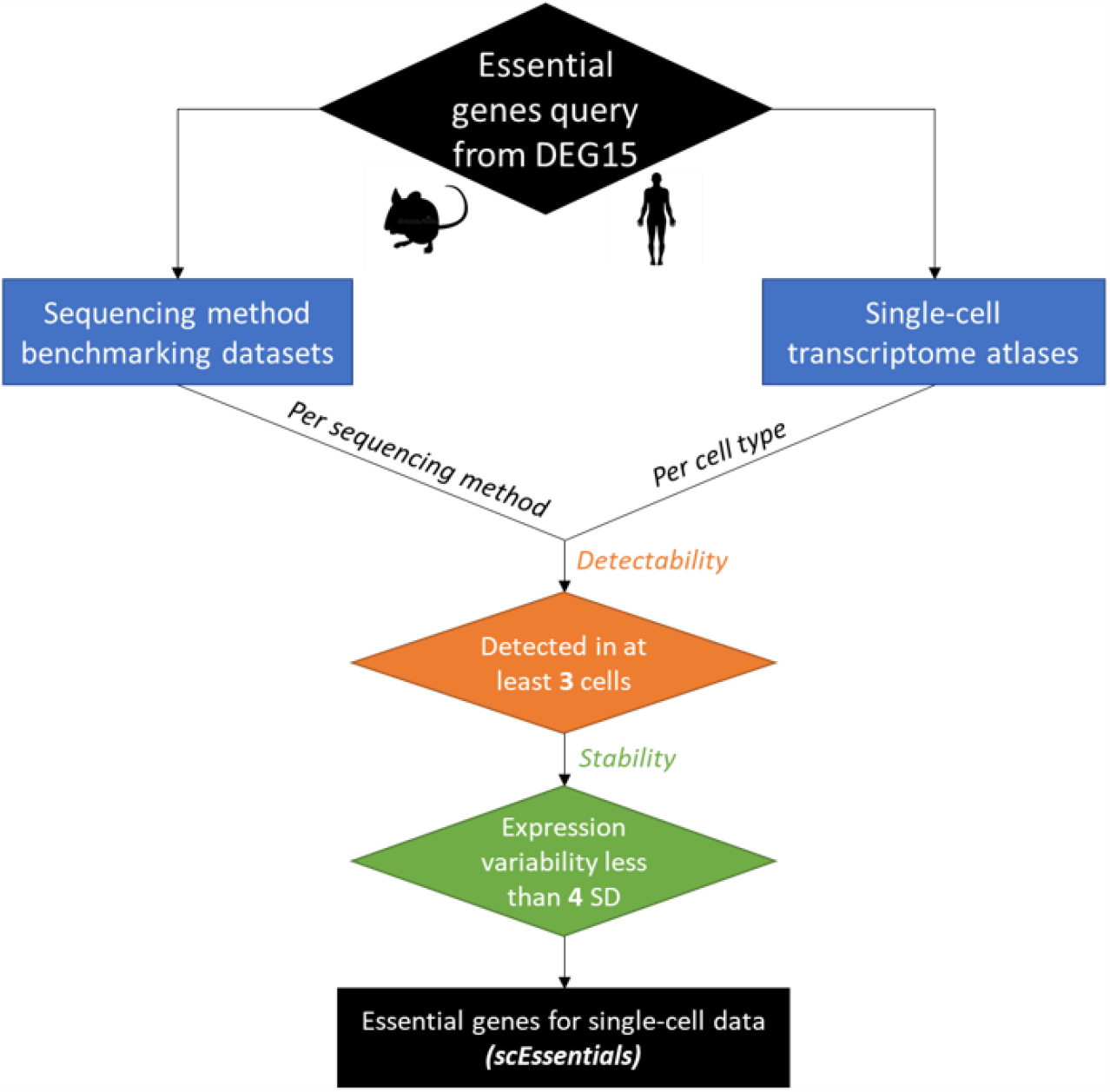
Overall workflow to determine essential genes for single-cell data (scEssentials) in mouse and human. Experimental-derived essential genes were queried from DEG 15 database (2) for mouse and human, respectively. For each species, datasets benchmarked popular-used sequencing methods and large-scale single-cell transcriptomic atlases were used to determine detectable and stable essential genes and denoted as scEssentials.

Additionally, we also we hypothesised that essential genes with excessive expression variations across different sequencing met were more likely to be affected by sequencing technology so that should be removed. To determine highly variable essential genes, we ranked essential genes based on their expressions in each sequencing method, where the highest values had the highest rank. As a result, any essential genes with average ranks across sequencing methods that were higher than four times of standard deviation of the distribution of the ranks were removed, remaining 1237 and 5770 essential genes for mouse and human, respectively.

### Application of scEssentials method to single cell tissue atlases

These essential genes were further assessed by over 50 cell types from TM and TS to confirm the essential genes detectability and stability. Similar to previous step, we removed the essential genes with low detectability (with less than 3 non-zero reads) and low stability (expression ranks higher than four times of standard deviation of the distribution of the ranks). In end, 733 and 1969 essential genes for mouse and human, respectively, were denoted as scEssentials (Supplementary File 1).

### Data normalisation, markers detection and stably expressed genes signatures

*SCTransform* (18) was applied to normalise the data by removing the batch effect for mESCs data and B lymphocyte data as well as regressing the mice/donor effect for TS and TM. Differential expression analysis was performed with the *MAST (19)* package in a 1-vs-all fashion within each tissue to identify cell-type-specific markers. The algorithm used a hurdle model to deal with the excessive zeros in the scRNA-seq data. The highly variable gene signatures were identified by *vst* method in *Seurat (v4*.*0)* (20) for each cell type.

The stably expressed genes and housekeeping genes were derived from three studies identified through statistical modelling and meta-analysis through large datasets (10, 21, 22). Only human data were available for three studies and hence used for our analysis. Specifically, only cytosolic ribosomal genes were selected from *Deeke et al* as these genes showed high expression and strong stability in bulk RNA-seq data. Gene lists used in the analysis were denoted as Supplementary Table 2.

### Constructing an Essentiality score

An *in silico* essentiality score was defined for each scEssential gene based on two main characteristics, non-cell-type-specificity and robustly high expression. Therefore, we leveraged the *PanglaoDB* (23) to quantify if any of the scEssentials have been identified as cell-type markers. Given a set of scEssentials, the raw essentiality score (ES) for gene *i* across *k* cell types is computed as:

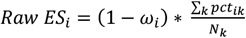

where *pct*_*ik*_ is the percentage of cells that expressed gene *i* in cell type *k*. Measuring the *pct* instead of mean expression for each gene provided a wider comparison range, while maintaining the characteristics of the high average expression given the high correlation between *pct*_*ik*_ and mean expression (Supplementary Figure 1). *ω*_*i*_ is the cell-type-specificity weight calculated from *PanglaoDB* as well as the frequency of gene *i* has been identified as a marker in TM and TS, namely,

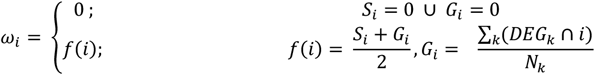

where *S*_*i*_ referred to sensitivity from *PanglaoGB*. Thus, genes that showed cell-type-specificity would be down-weighted while those with a higher number of cells expressed would be up-weighted. Notably, the distribution of raw ES was right-skewed (Supplementary Figure 2), with median ES score at 20. To mitigate the skewness and achieve compatible values, logarithm_2_ transformation was applied on raw ES.

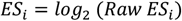

**Figure 2.**
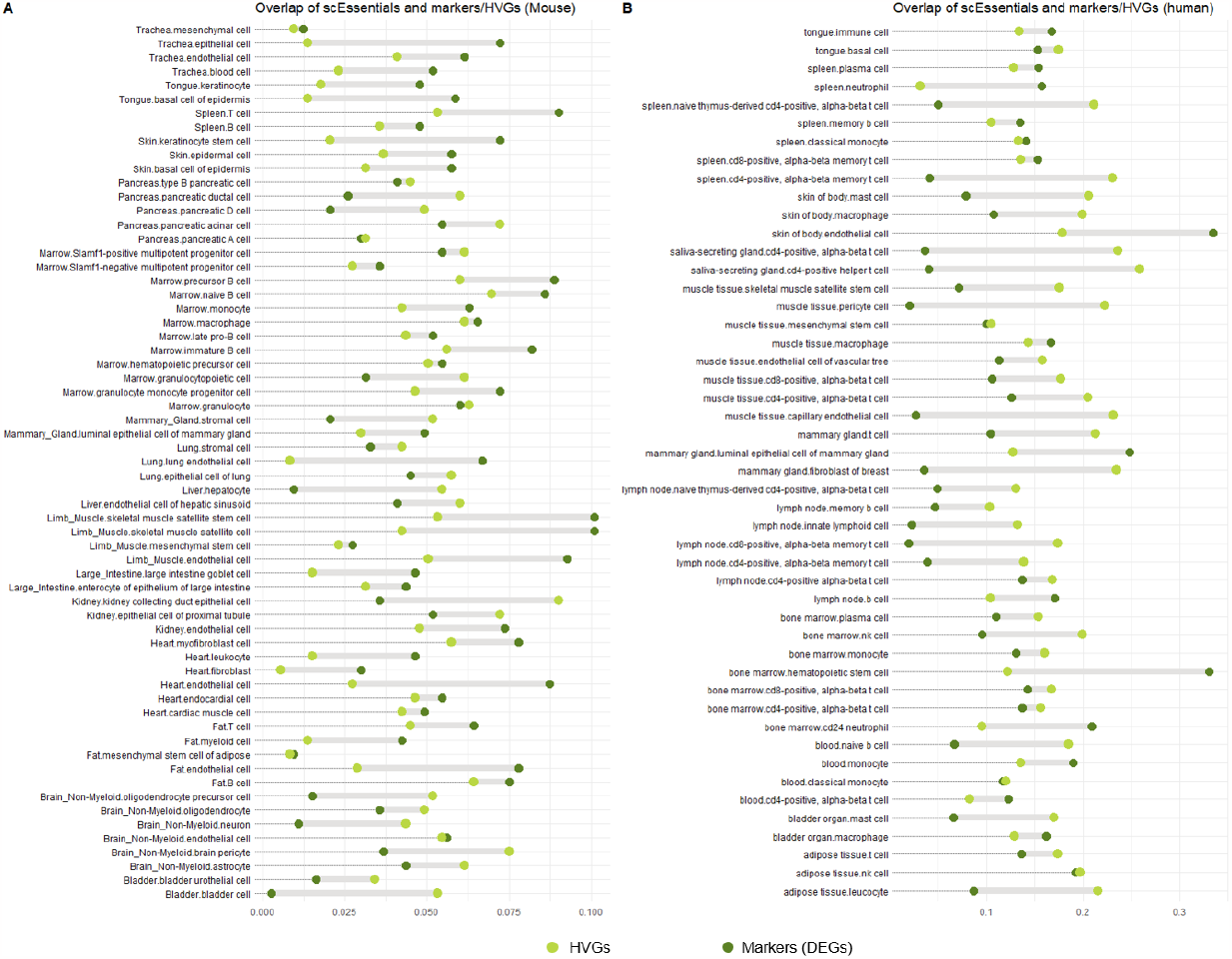
Overlap of scEssentials with cell-type specific differentially expressed markers (DEGs) and highly variable genes (HVGs) across cell types in A) Tabula Muris (mouse) and B) Tabula Sapiens (human). Cell-type markers were identified by the hurdle model from *MAST* (19) and HVGs was referred to as the HVGs identified by variance stabilising transformation, with the same number of scEssentials. Dark green represented the overlap with DEGs and light green represented the overlap with HVGs.

### Cell type similarity based on essential genes and visualisation

Pearson correlation was applied to measure the relationships among sequencing methods as well as the cell types from TM and TS based on the scEssentials genes’ expression. The correlation plots that illustrated the similarities of different aspects under essential genes were constructed by *corrplot (v0*.*9)* (24).

### Measure gene expression variability in scRNA-seq data

To investigate and compare gene expression variability between scEssentials and non-scEssentials at different molecular levels, we used *scran* (*v1*.*20*.*1*) to measure the cell-to-cell variability with mouse/donor ID as covariate in the regression model. As suggested in our previous comparative study (25), *scran* enables accurate calculation of cell-to-cell variability in scRNA-seq data.

### Assessing significance of the essentiality score relative to a random gene set

To characterise the scEssentials, we used random sampling gene lists as the control to compare against. All the random gene list was sampled under the full data matrix with the same number of essential genes and the random sampling was computed 10 times.

### Gene pairs correlation level and co-expression networks based on scEssential at the cell-type level

A modified Spearman correlation analysis was performed to identify the correlation coefficients between gene pairs in scRNA-seq data by *correlatePairs* from scran (26). As excessive zero would mislead the correlation, we excluded the genes expressed in less than 10% of each cell type population. After BH correction, *FDR<0*.*05* was used to determine significant gene pairs. The proportion test was applied to determine the significant difference in the number of gene pairs. Given the number of significantly correlated gene pairs was substantially different across random gene lists, we only extracted the top 100 most correlated gene pairs (both positively and inversely correlated) from each cell type or the maximum number if the total number is greater than 10 and less than 100, otherwise the cell type will be removed. The *Kruskal*-*Wallis test* was applied to determine the level of significant gene pairs’ correlation coefficients to compare scEssentials and random gene lists for each cell type. Cell type To compute the correlated scEssentials gene pairs, boxplots for the absolute r values and histogram representing the mean absolute correlation coefficients for each cell type were combined by *ggpubr (v0*.*4*.*0)* package (27).

Co-expression networks were constructed for scEssentials and random gene sets based on significantly correlated gene pairs *(P-value < 0*.*01)*. Betweenness and normalised node degree of each network were calculated by *igraph (v1*.*2*.*6)* (28). Eventually, a set of robustly correlated gene pairs was aggregated if they were significantly correlated in more than half of the total cell types and the network was plotted by *igraph*. A fast greedy modularity optimization algorithm was applied to the shared network to identify communities and related functions were computed against the GO database.

### Gene sets from curated pathway databases

To measure the proportions of essential genes in the curated pathways, we leveraged the Hallmark and KEGG pathways gene lists obtained from *msigdbr (v 7*.*4*.*1*.) package (29). The number of essential genes found in each pathway was collectively compared to the random gene list. The *Kruskal*-*Wallis test* was applied to measure whether the number of the pathways genes were more in the scEssentials genes differs from 10 other random gene lists.

### Gene damage index (GDI)

The *in silico* GDI score was estimated for each gene’s mutation damage that has accumulated in the general population under a comprehensive multi-level analysis, which helps to predict whether a gene is likely to harbour disease-causing mutation (30). A high GDI score represents the frequently mutated genes that are more likely to cause inherited and rare diseases but unlikely to make a disproportionate consequence. Whereas, with a low GDI score, the rarely mutated genes, are more likely to be disease-causing. The *Kruskal*-*Wallis test* was applied to test the Phred-scaled GDI indexes between scEssentials genes and random gene lists. We applied Spearman correlation to measure the relationship between normalised GDI score and scEssential genes.

Additionally, scEssentials expression variability differences in three categories low GDI (91 genes), medium GDI (1728 genes) and high GDI (39 genes) were compared using the *Kruskal-Wallis* test to calculate the significance. To mitigate the impact of unequal sample sizes, we downsampled the number of genes in the medium GDI to 39 through *downsample* method in *caret (v6*.*0-90)* (31). This procedure was repeated 10 times.

### Chromatin Accessibility level inferred from single-cell Assay for Transposase-Accessible Chromatin sequencing (scATAC-seq)

To investigate the chromatin accessibility of the scEssentials genes, we utilised cell types in marrow tissue that existed in both TM and mouse scATAC-seq atlas (32). Consisting with the cell-type selection in the TM dataset, we only included five cell types that have over 100 cells including hematopoietic progenitors, B cells, macrophages, monocytes and immature B cells. The processed binary count matrix was downloaded from the repository, https://atlas.gs.washington.edu/mouse-atac/data/ and the annotation followed the original paper description. Only the promoter transcription starting sites that overlapped the processed peaks with were included. Spearman correlation was applied to measure the relationship between the chromatin accessibility level and the essentiality score for genes within each cell type.

### Transcription factors (TF) identification

*AnimalTFDB 3*.*0* was used to search for mouse TFs in the scEssentials (33). To calculate the overlap enrichment of TFs in the scEssentials genes, we applied the hypergeometric test with all protein-coding genes as the background genes. In total, 24,351 mouse protein-coding genes were extracted from the Mouse Genome Informatics database (34). *Phyper* in R was applied to calculate the significance.

### Modelling the scEssential gene expression changes in ageing

Only mice from 3-month, 18-month and 24-month that have been sequenced by FACS-smartseq2 methods from Tabula-Muris-Senis (35) were included in the analysis to reduce the data sparsity. We excluded the cell types that had less than 100 cells. To understand how essential genes change during ageing under varieties of cell types, we performed a generalised linear regression model on each gene separately by analysing the age and cell type interaction while controlling sex as covariates. For each gene *i*, the model is

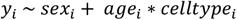

where *sex*_*i*_ (male and female) was corrected in the model and interaction term *age*_*i*_ * *celltype*_*i*_ measured the effect of age groups (3-month, 18-month and 24-month) depending on different cell types for each gene *i*. P-values were corrected by BH methods. Genes that showed significant changes across age groups investigated the expression changes under the effect of various cell types. Genes that showed no significant difference in age groups further deciphered the interactions with different cell types. Barplots and boxplots were generated to demonstrate the performance of scEssentials under age covariates and their effect on various cell types.

### Pathway over-representation analysis

The pathway over-representation analysis against the GO database was performed on the scEssentials genes that significantly changed during ageing via *clusterProfiler (v4*.*0*.*5)*, with genes detected from TM and TS as universe gene sets (36). The top 10 significant enriched pathways based on p-values were plotted.

### Statistical analysis and visualisation of data

All the random gene list was sampled under the full data matrix with the same number of essential genes and the random sampling was computed 10 times. Plots were generated through *ggplot2* unless stated with the seed. The BH adjustment method was applied and the significant adjusted p-value level was set to 0.05. The code to generate the plots and processing scheme can be found at the GitHub repository https://github.com/huiwenzh/scEssentials

## Results

### Constructing the landscape of essential genes at single-cell level (scEssentials) using Tabula Muris and Tabula Sapiens

Essential genes have been tested on multiple cell lines and individuals and demonstrate their critical roles in cell survival. In fact, not all genes are expressed in diverse cell types so whether these essential genes are essential for all cell types and their expression patterns remain unclear. Therefore, it is critical to investigate the cell type-specific expression of essential genes across all tissues in a species-specific manner as the differences between human orthologs of mouse essential genes and human essential genes are inevitable (11). In this study, we used datasets that benchmarked popular-used sequencing methods as well as two large-scale single-cell atlases, Tabula-Muris (TM) and Tabula-Sapiens (TS), to investigate the characteristics of essential genes at cell type-level in mouse and human, and determined a set of essential genes that were generalisable to different scRNA-seq data (scEssentials).

We considered the detectability and stability in the essential genes to ensure that scEssentials were applicable and reliable (Figure 1, more details in Method). The detectability of essential genes were measured whether they can be identified in more than 3 cells per sequencing method or per cell type in the single-cell atlases. On average 80% of essential genes were detected across methods for both mouse and human datasets. Especially for non-UMI methods, more than 90% of essential genes can be detected (Supplementary Figure 3). Conversely, decreased detectability was found in TM (70%) across 68 cell types and substantially more in TS (64%) across 53 cell types, suggesting the sequencing variations on the heterogeneous biological samples. Additionally, we removed highly variable essential genes either due to the technical (different sequencing methods) or biological (different cell types). Interestingly, we observed higher variations in essential genes from technical aspect rather than biological (Supplementary Figure 3-4). In end, we retained 733 and 1969 essential genes for mouse and human, respectively and noted as scEssentials (Supplementary File 4.1). The correlation plots based on the scEssentials expression demonstrated high correlation among various cell types in both TM (*mean r* = 0.75) and TS (*mean r* = 0.80) (Supplementary Figure 5A-B), where each tissue-cell-type was closely clustered with the same cell types rather than the tissue type.

**Figure 3.**
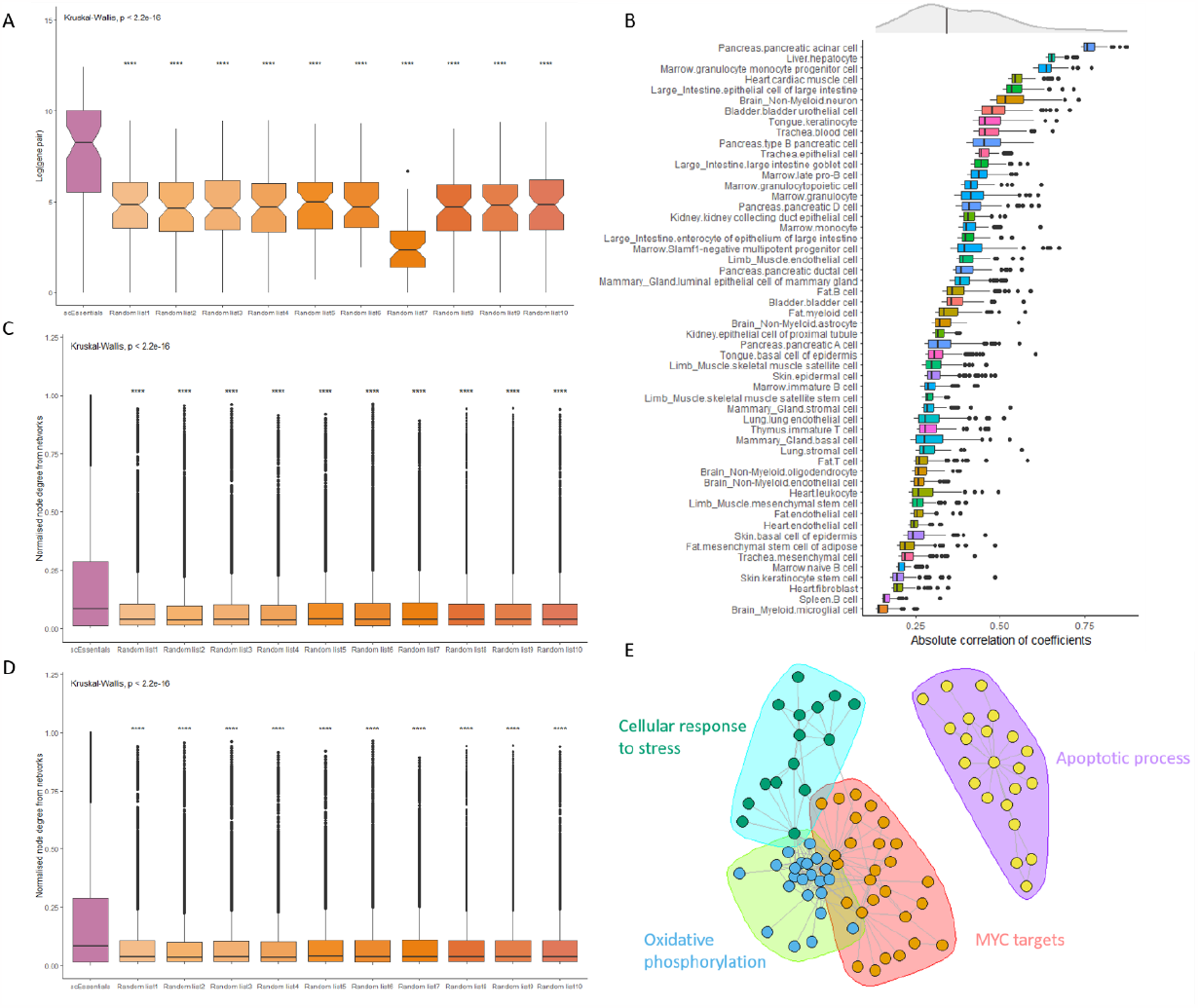
Analysing correlation on scEssentials gene pairs and 10 random gene lists. Boxplots illustrated A) the number of significant gene pairs (*FDR P-value* <0.01); B) top 100 most correlated scEssentials gene pairs across cell types (coloured by tissue origin); C) the network betweenness; D) the network normalised node degrees that constructed from scEssentials and random gene lists. The Kruskal-Wallis test applied to determine the significance (* p<0.05; ** p<0.01; *** p<0.001; **** p<0.0001). E) A core network from significantly consistent correlated scEssentials gene pairs. The communities were identified by a fast greedy modularity optimization algorithm.

**Figure 4.**
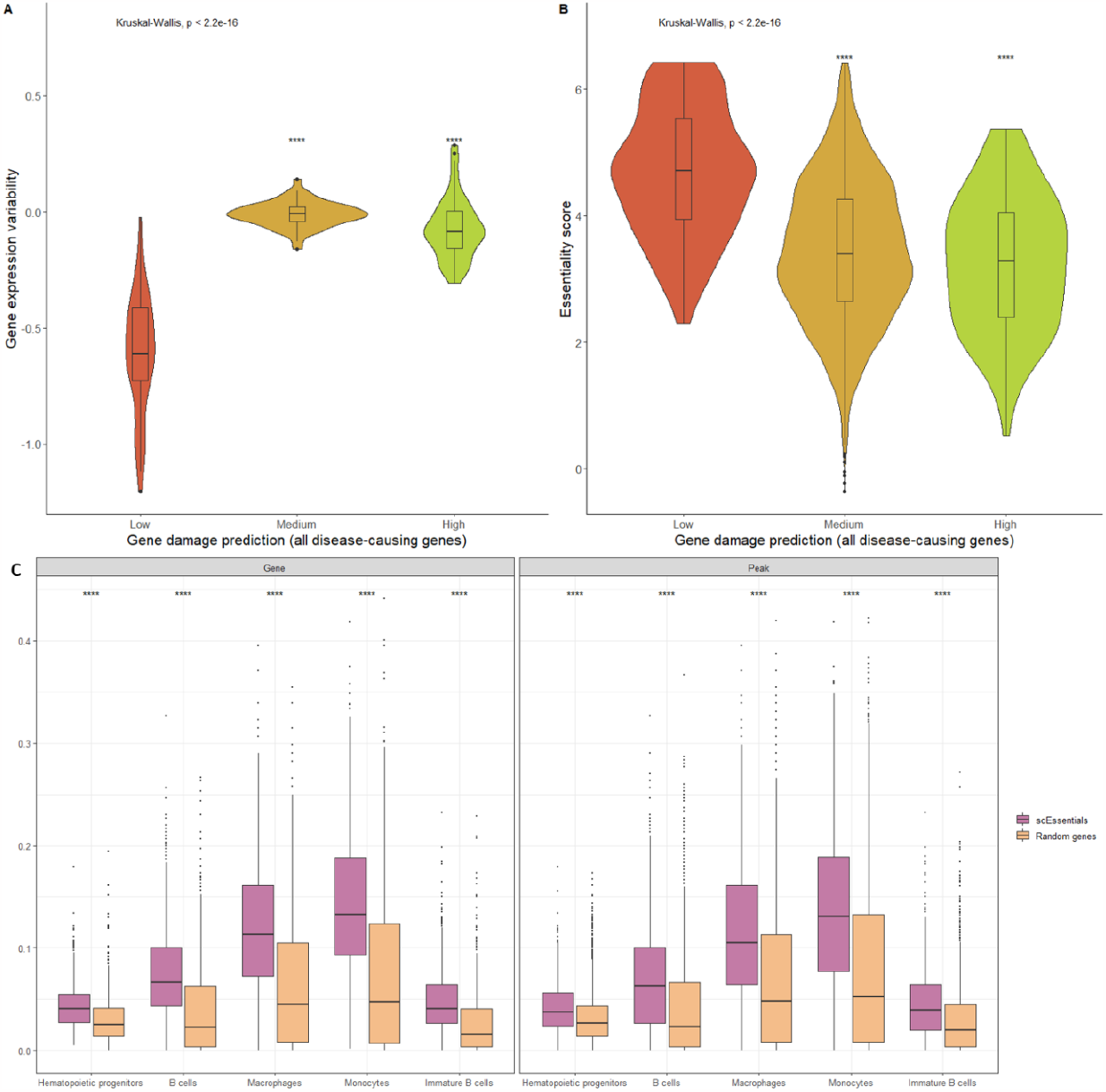
Analyses of mouse scEssentials genes with gene mutation index and chromatin accessibility level. A) Violin plot showed the gene expression variability level within low, medium and high risks groups. B) Boxplot showed the ES score difference among low, medium and high risks groups. Kruskal-Wallis test was applied to determine the significance. C) Chromatin accessibility level for five cell types from scATAC-seq atlas in scEssentials genes and a random gene list. The accessibility was measured by peaks and gene level. Wilcoxon ranked sum test was applied to determine the significance (* p<0.05; ** p<0.01; *** p<0.001, **** p<0.0001).

**Figure 5.**
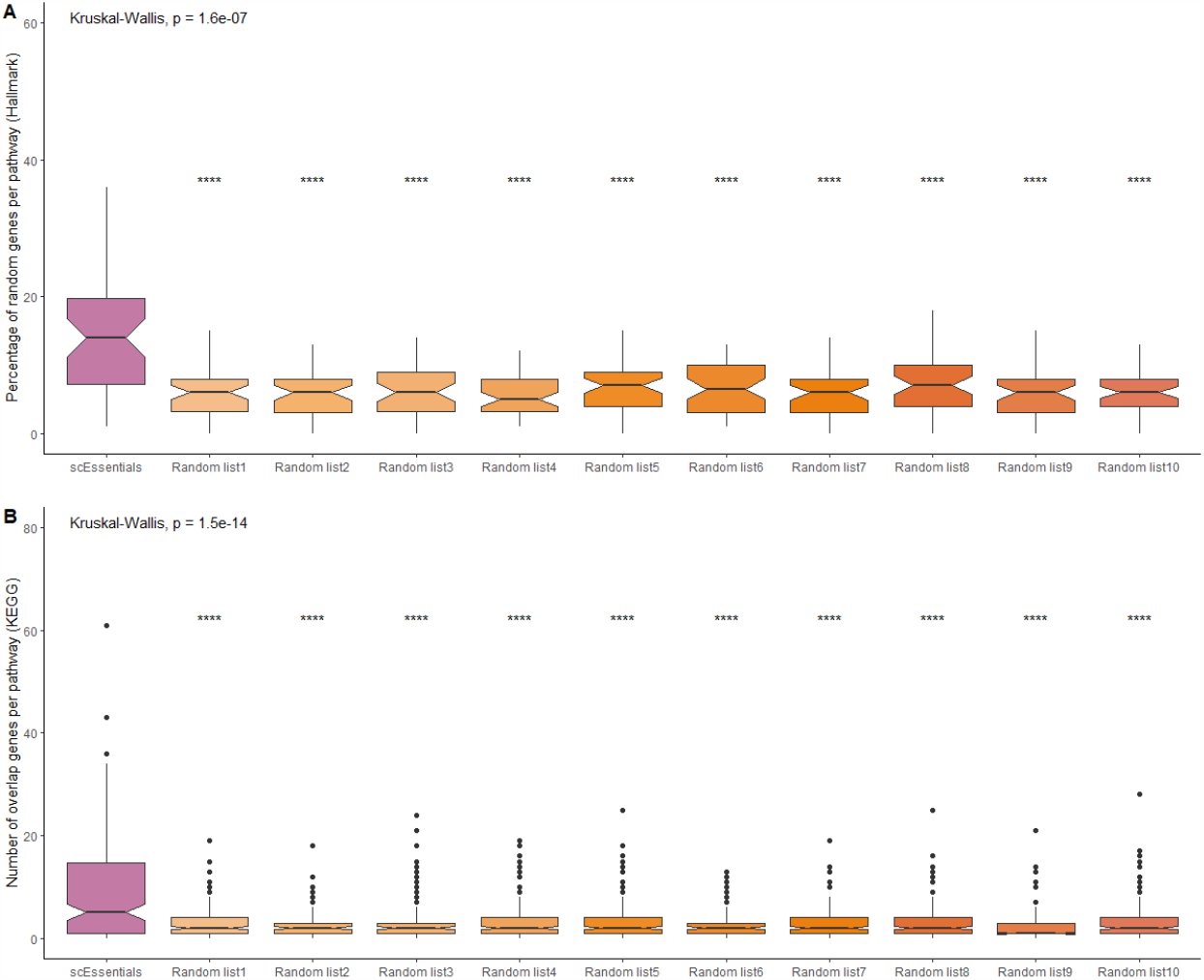
Percentage overlap with two pathway databases A) Hallmark database and B) KEGG. In total, there are 50 pathways in Hallmark database and 186 pathways in KEGG database. scEssentials genes and 10 randomly sampled gene lists were plotted in the boxplot and the *Kruskal-Wallis* test was applied to determine the significance. * p<0.05; ** p<0.01; *** p<0.001, **** p<0.0001.

### scEssential genes are highly expressed in all cell types with minimal overlap with cell-type specific markers

One of the key aspects of essential genes is the relatively higher expression compared to other genes (5). We confirmed this phenomenon at the cell-type level as scEssentials showed significantly higher expression among all cell types in both TM and TS data as compared to random gene lists (*Wilcoxon ranked sum test, p<0*.*05*) (Supplementary Figure 6). This supports the observation that scEssentials had higher expression values based on tissue-level RNA-seq data (5). Additionally, scEssentials retained a higher percentage of cells that have expressed the genes among all cell types (*Wilcoxon ranked sum test, p< 0*.*05*) (Supplementary Figure 7).

**Figure 6.**
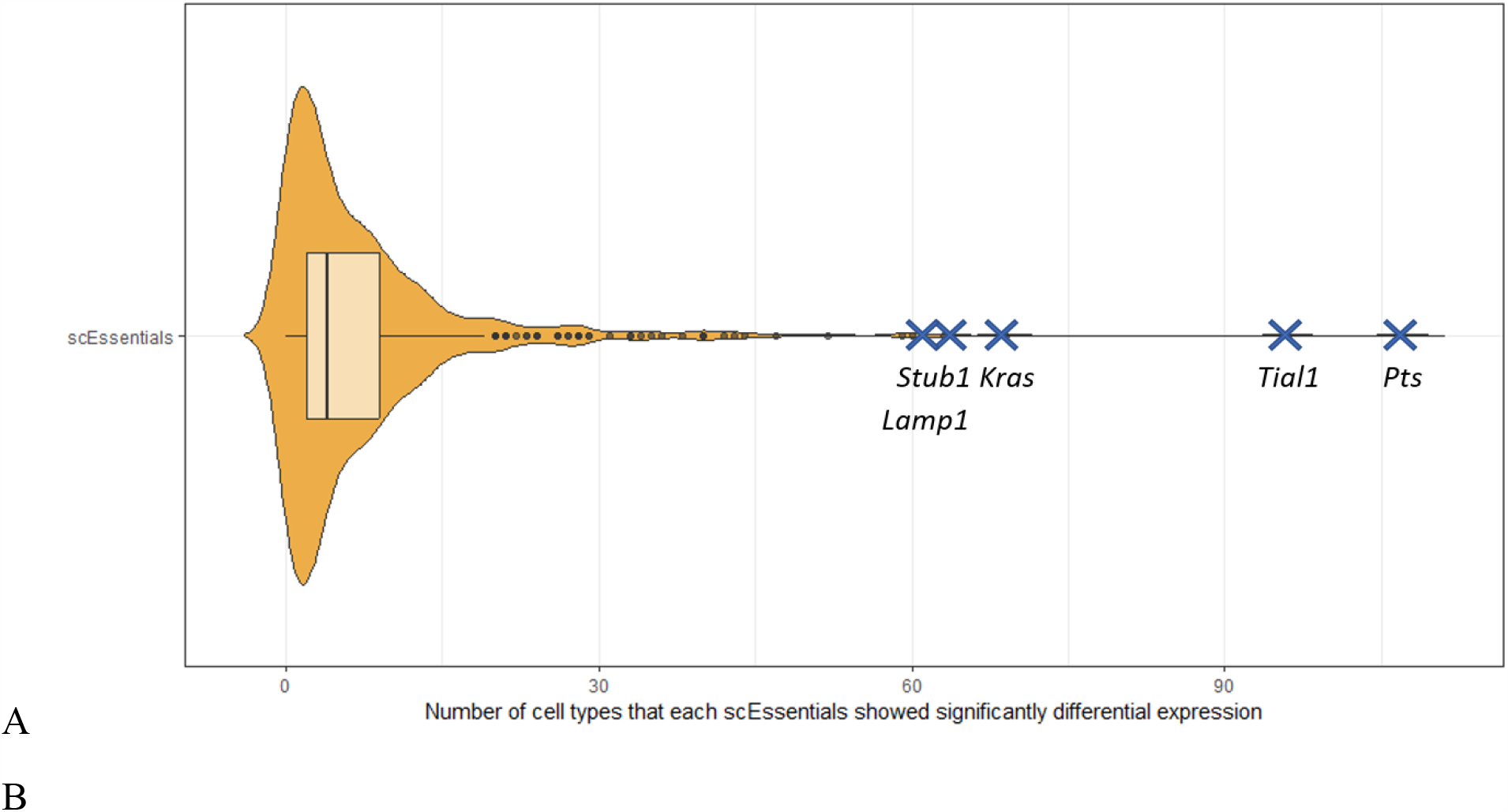
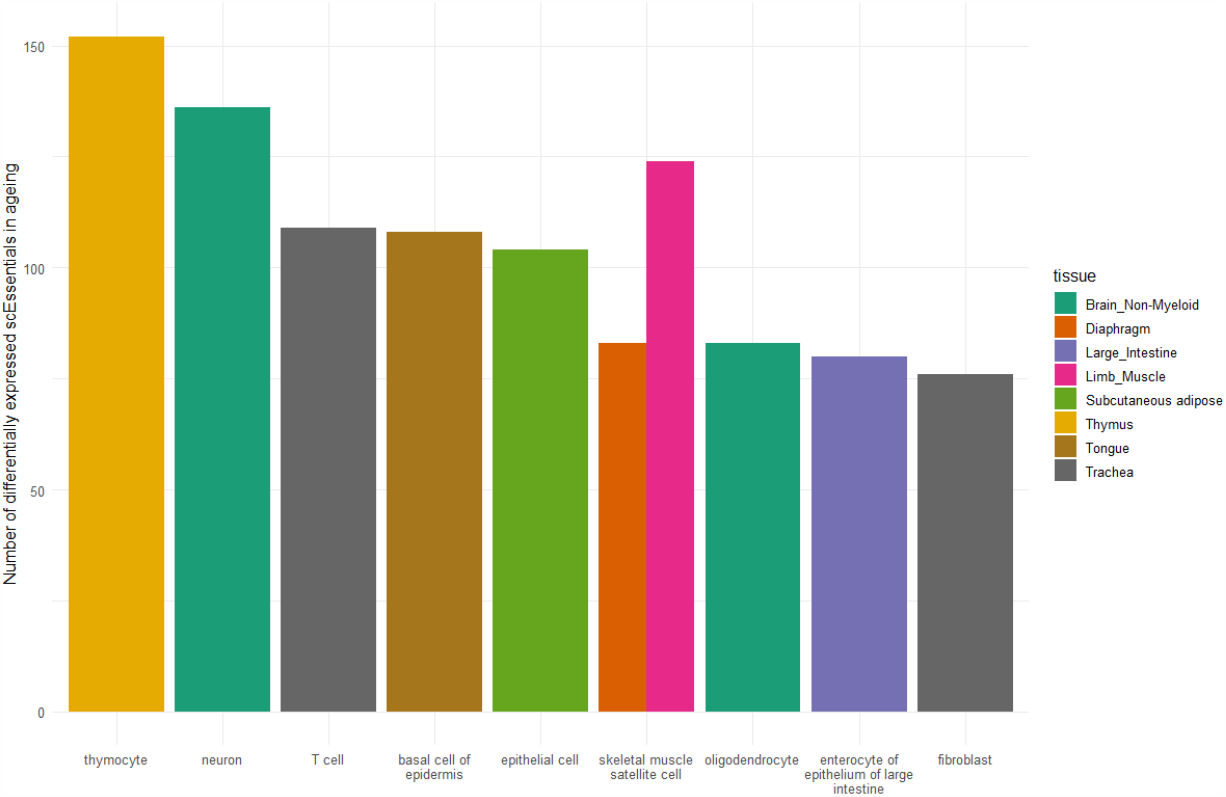
Dysregulation of scEssentials genes among cell types in ageing. A) Density plot for each scEssentials genes that were significantly differentially expressed. Genes with the top number of significant changes among cell types are highlighted. B) Bar plot illustrating the top 10 cell types with the highest number of scEssentials gene that showed significant change in ageing. Each bar was coloured by the tissue it is originated from.

A key consideration that arises when analysing scRNA-seq data is the identification of cell-type-specific markers that include differentially expressed genes (DEG) and highly variable genes (HVG), and these gene sets are used to improve the assignment of cell identities. We utilised a hurdle model to determine the makers and then examined whether there was an overlap with the genes identified by scEssentials to see if they were differentially expressed in any cell types (20). As expected, most of the cell-type-specific DEGs shared minimum overlap with the scEssentials for both mouse and human datasets (median 5% in TM; median 11% in TS). In addition, we also compared genes identified by scEssentials with a cell type marker database *PanglaoDB (23)* to further measure cell-type specificity based on external evidence. Although we found 119 mouse and 175 human scEssentials were identified to be cell-type markers, with only about one-sixth of cell types in the *PanglaoDB* were overlapped with cell type in TS. Similarly, the HVGs and scEssentials with TM data showed an even smaller overlap (median 4%) (Figure 2). However, the HVGs shared a higher proportion with scEssentials with TS data (median 27%) (Figure 2), which might result from the type of treatment or disease occurred in the donors. Overall, the increased number of scEssentials in common with DEGs and HVGs in TS, reflecting the increased complexity and heterogeneity in human data compared to mouse data.

In contrast to HVGs, ribosomal genes, stably expressed genes (SEGs) and housekeeping genes were also compared to scEssentials because of the related underlying characteristics. In general, most of the SEGs and housekeeping genes lists were curated by different statistical algorithms or meta-analyses of related literature which often showed inconsistency and method-dependency. By comparing three gene lists with scEssentials, we observed an overlap of around 56% of total gene lists regardless of the size of different gene lists (Supplementary Figure 8). Interestingly, the majority of ribosomal genes (82%) overlapped with at least one gene list, as they posse high expression levels and critical roles in cell growth and proliferation (37). In addition, scEssentials showed the highest overlaps with housekeeping genes and these overlapped genes were involved in RNA catabolic process and splicing functions. Overall, scEssentials demonstrated high expressions and minimum overlaps with markers identified from DE methods and marker database in both TM and TS. Furthermore, scEssentials also showed variations in some cell types (for example, muscle satellite cells) in TM and increased levels of variations across all cell types in TS while retaining high overlaps with other known gene lists with high stability.

Here, we developed an *in silico* essentiality score (ES) for scEssentials that incorporated the detectability and gene marker sensitivity (see details in Method). Any scEssentials held no evidence being any cell-type-specific markers and with high detectability across cell types would be up-weighted so that higher ES suggested greater importance that scEssentials had. Particularly, the top-ranked genes were mostly ribosomal genes as well as apoptosis genes for both human and mouse datasets, which agreed with the previous study that summarised essential genes’ potential categories (5) (Supplementary File 2).

### scEssential gene pairs are highly and consistently correlated in various cell types

To explore the relationships and interactions of genes identified by scEssentials, we investigated the gene pair co-expression as compared to other genes. As scEssentials showed a higher frequency among biological pathways, we hypothesised that scEssentials have closer relationships to maintain cell revival and function as compared to random genes. To capture this relationship, traditional correlation analyses like Pearson and Spearman are commonly used. However, these methods are often biased by excessive zeros resulting in unreliable correlation coefficients (38). Therefore, the modified Spearman correlation analysis based on the permutation test was applied to determine the gene pairs’ correlation (26). We firstly measured the correlation of all the genes in 10 randomly sampled gene lists with the same size of scEssentials as well as scEssentials (see Method). The boxplot demonstrated an increased average number of significantly co-expressed gene pairs in scEssentials as compared to the randomly sampled gene lists (Supplementary Figure 9A, *Kruskal*-*Wallis test, P-value < 0*.*05*). When inspecting the correlation coefficient values, we observed suspicious perfect correlation coefficients (*Rho = 1*) in the random gene pairs. By further examining these perfect correlations, we found that these gene expressions were predominant zeros (more than 90% zero values) and the perfect correlations resulted from just 1-2 cells’ expression values. Therefore, to reduce the false discovery rate, we only included the genes that were expressed in more than 10% of the cell type sizes for the comparison. Consequently, we observed a substantial increase in the number of refined correlated gene pairs with scEssentials compared to all random gene lists (*Kruskal*-*Wallis test, P-value < 0*.*05*) (Figure 3A), with about a 30-fold difference. Interestingly, even though the sizes were dramatically different, the number of significant co-expressed gene pairs was proportional between scEssentials and randomly sampled gene lists. It indicated the levels of tight co-regulation were naturally distinct across different cell regardless of the genes used for the measurement. For each cell type, scEssentials obtained a consistently higher number of significantly correlated gene pairs for 84% of cell types (Supplementary Excel File 3, 57/68; *proportion test, P-value < 0*.*05*). Interestingly, cell types that did not show higher number of correlated gene pairs in scEssentials mostly consisted of precursor cells from different tissue origins and cell types with relatively smaller sample sizes (lower than median sample sizes across all cell types).

Furthermore, we aimed to examine the correlation coefficient level in the scEssentials as compared to randomly sampled gene lists to assess how strong the co-expression are. Given the significant difference in the number of refined correlated gene pairs between scEssentials and random gene lists, we extracted the top 100 most correlated gene pairs from each gene list and compared the coefficients (see Methods). Consistently, we observed significantly increased correlation coefficients in scEssentials as compared to other 10 random gene lists in the all cell types, except for two progenitor cells in marrow tissue (Supplementary Figure 9B-C, 66*/68; Kruskal*-*Wallis test, P-value < 0*.*05)*. Furthermore, we investigated the degree of scEssentials gene pairs’ correlation across cell types. The absolute coefficients for the selected correlated scEssentials gene pairs across various cell types were shown in Figure 3B, with the mean absolute coefficient value at 0.36. Unlike the consistent expression pattern for individual scEssentials levels, the correlation range between scEssentials varied among cell types. Interestingly, the most tightly correlated cell types were those specialised epithelial cells, for example, pancreatic acinar cells, hepatocytes and cardiac muscle cells. Overall, scEssentials correlation coefficients showed consistently high values and higher cell-type variations as compared to a single gene level.

Finally, we constructed the co-expression network from the common significantly co-expressed scEssentials gene pairs across cell types to assess the functional relationship with respect to cell maintenance.

Betweenness and normalised node degree were calculated for each network to measure how many shortest paths pass through a node and how connected each of the nodes in the network is. Higher betweenness and normalised node degree levels demonstrated a higher degree of network density. Ultimately, on average networks constructed by scEssentials genes across cell types showed significantly higher node degrees and the nodes (genes) in each network showed higher betweenness as compared to random gene lists, which indicated that the scEssentials correlation network tended to be more closely interact to maintain functional roles (Figure 3C-D). Furthermore, we identified a core co-expression network that constructed from common co-expressed gene pairs in more than 30 cell types, which robustly connected among various cell types, including 86 nodes (genes) and 227 vertices (correlations). Five community structures were identified by a fast greedy modularity optimization algorithm and each community was functionally related to apoptotic, oxidative phosphorylation and cell maintenance (Figure 3E).

### scEssential genes displayed a low frequency of somatic mutation that is associated with the low level of gene expression variability

Given the stability that scEssentials possess, it is unsurprising that such stability is present in other molecular levels like the stable encoded protein (5). To further investigate scEssentials characteristics at other molecular levels, such as genetic mutations, the gene damage index (GDI) that quantified the mutation frequency of protein-coding genes in the general population was compared (see Methods) (30). Consequently, scEssentials showed a significant correlation with low gene damage prediction (all disease-causing genes) and decreased normalised GDI value as compared to other protein-coding genes (*Chi-square* test, *P-value < 0*.*05*) (Supplementary Table 2). Lower GDI indicated the less frequently mutated genes which are more likely to be disease-causing, representing the critical roles of scEssentials at the genetic level to maintain regular cellular activity.

Next, we interrogated whether such mutation frequency is associated with gene expression variability. These GDI values were categorised into low, medium and high risk levels based on different thresholds (30). To accurately measure scEssentials cell-to-cell variability in scRNA-seq data, we used *scran* on each cell type as suggested best performance in Chapter 3 (25, 39). Interestingly, significantly low gene expression variability was found in the low GDI group on average which aligns with the interpretation that genes with increased expression stability are associated with a lower risk of mutation frequency in disease-causing genes. Although there was no significant mean difference in the gene expression for the medium and high GDI groups, the high GDI group demonstrated significantly increased levels of cell-to-cell variability (Figure 4A). Specifically, we observed a commonly increased level of gene expression variability in the genes with medium damage and high damage as compared to low damage for over 42% of cell types (*22/53; Kruskal*-*Wallis test, FDR P-value <0*.*05*) (Supplementary Figure 10A). Additionally, genes in low GDI group showed an increased number of more stable genes (negative values) than more variable genes (positive values) as compared to other groups. The increased gene expression variability illustrated the potential impact of somatic mutation burden on the stochastic heterogeneous gene expression. We also found that most of the stable gene expression resulted from a set of ribosomal genes whereas the top variable genes were associated with the regulation of cell death (*Msigdb*). However, when comparing the scEssentials with high GDI to non-scEssentials in high GDI group, scEssentials in high GDI showed a significantly reduced level of gene expression variability in more than 77% of cell types (41/53; *Wilcoxon ranked sum test, FDR P-value <0*.*05*) (Supplementary Figure 10B). It emphasised the stability of the scEssentials expression level was robust even though they possess high risks for damage from mutations in disease.

Lastly, we analysed the relationship between ES and the normalised GDI score to assess the ability of ES on capturing the mutation frequency (See Methods). As a result, low GDI scEssentials found significantly high ES as compared to medium and high GDI groups (*Kruskal*-*Wallis test, P-value < 0*.*05*; Figure 4). Furthermore, ES values were significantly inverse correlated with the normalised GDI score across all cell types, with median correlation coefficient at -0.4 (*Spearman correlation; P-value < 0*.*05;* Supplementary Figure 10C), suggesting the mutation frequency can be well-captured by ES to reflect the biological stability in scEssentials. Hence, low frequency of somatic mutation and low level of gene expression variability in scEssentials were supported by these analyses and effectively captured by ES values.

### scEssentials genes showed significant higher level of chromatin accessibility and number of transcription factors

In addition to gene mutations, gene expression is regulated by epigenetics factors such as chromatin packaging and accessibility. The previous study has reported that essential genes have been shown as hubs of active histone modification processes and DNA methylation patterns (5). Recent advances in single-cell Assay of Transposase Accessible Chromatin sequencing (scATAC-seq) enable cell-type level investigation of the regulatory landscape without prior knowledge of regulatory elements. To investigate the scEssentials genes’ active regulation, we used a large mouse scATAC-seq atlas that captured chromatin availability from multiple tissues/cell types in 17 young mouse (32). Bone marrow was selected for this comparison as it shared the most number of cell types between scRNA-seq and scATAC-seq data, resulting in five cell types with 100 cells (see Methods) (Supplementary Figure 11A). Compared to the random gene list, the peaks that intersected with the transcription starting site (TSS) of scEssentials genes showed significantly higher levels of accessibility (*Wilcoxon ranked sum test, P-value < 0*.*05*) (Figure 4C). Furthermore, we aggregated multiple peaks that were annotated to one gene to reduce the variation. At the gene level, scEssentials showed further increased degrees of chromatin accessibility level compared to the random gene lists at peak level (*Wilcoxon ranked sum test, P-value < 0*.*05*) (Figure 4D).

To evaluate whether the ES was able to identify trends in the level of chromatin accessibility inferred from scATAC-seq data, we computed the correlation between ES and chromatin accessibility for five cell types separately. As expected, ES significantly correlated with the level of accessibility in every cell type (*Spearman correlation, median r=0*.*21; P-value < 0*.*05*) (Supplementary Figure 11B), suggesting the robustness of the ES in capturing active regulatory inferred from chromatin accessibility level. Furthermore, scEssentials had a significantly higher number of transcription factors (TFs) that are responsible for controlling gene expression (*Hypergeometric test, P-value < 0*.*05;* Supplementary Figure 11C) (see Methods). Taken together, we demonstrated that scEssentials possessed tight regulation at the epigenetics level to maintain stable gene expression and the corresponding ES values were robust to capture the gene importance at different molecular levels such as genetics and epigenetics.

### scEssential genes showed high involvement in the curated pathway databases

Biological pathways are comprised of sets of interactions between genes that produce a particular product in a cell. Therefore, genes in the pathways are more likely to modify the cellular functions than the genes that are not in the pathway. Kyoto Encyclopedia of Genes and Genomes (KEGG) database is one of the most widely used resources for understanding biological systems (40) and often viewed as the gold standard for pathway annotation. More recently, the hallmark database has effectively summarised the pathway information by reducing gene set redundancy (41). By leveraging the comprehensive and concise nature of both pathway databases, we investigated the involvement of scEssentials in more than 200 biological pathways. The involvement of scEssentials in these pathways against randomly sampled gene lists under the same size was measured by assessing the degree of overlap between the gene lists and each pathway to demonstrate the importance of scEssentials with respect to any random genes. For each pathway from both databases, we observed a significantly increased contribution of scEssentials as compared to 10 randomly sampled gene lists (*Kruskal-Wallis test, P-value < 0*.*05*), indicating the fundamental roles of scEssentials played in the various functional pathways to maintain biological functions (Figure 5). Interestingly, although overlaps of scEssentials in the pathways increased with larger pathway sizes, higher variation was found in pathways with sizes greater than 100 (Supplementary Figure 12). For instance, large pathways with the highest number of scEssentials overlap in the Hallmark database was mTORC1 signalling (*N=201, overlap=36*) while the lowest overlap was shown in gene lists related to angiogenesis (*N=36, overlap=1*). Notably, essential genes were mostly identified by experimental functional screening or knockdown experiments with well-characterised genes (42), which may exist an over-annotation effect in the curated databases. However, the usage of 10 randomly sampled gene lists in the analyses maximally reduced the effect and supported the evidence that scEssentials functioned as a core component among various biological pathways.

### Dysregulation of scEssentials genes showed cell-type-specificity in ageing

Transcriptional dysregulation is commonly observed in ageing. As numerous studies have described ageing tissue-specificity and cell-type-specificity across organisms (43-46), we then investigated whether the scEssentials genes would change during ageing given the stability and essentiality characteristics of these genes. To examine this hypothesis, we applied a generalised linear regression model to over 130 cell types to investigate the interaction between cell types and age groups for the scEssentials from Tabula-Muris-Senis (TMS) database (see Methods). For each gene, we examined covariate age as well as the interaction of age groups for each cell type. Surprisingly, we found that only 22% (162/733) of scEssentials genes’ expression dysregulated with age covariate, indicating the importance of the scEssentials regardless of ageing phenotypes. Additionally, these genes were found to be significantly associated with DNA repair and apoptotic pathways (Supplementary Figure 13A). These pathways are the signalling cascades triggered by DNA damage, which is the primary cause of ageing (47). We further compared the ES score between the significantly dysregulated scEssentials genes under age covariate and other non-significant ones. Interestingly, we did not observe a significant difference in ES score for the genes that had associated with age with respect to the genes that did not change with age (*Wilcoxon ranked sum test, P-value = 0*.*08*) (Supplementary Figure 13B).

Additionally, we explored the modification of scEssentials at the cell type level for the remaining scEssentials (non-significantly changed with age covariate). Inspecting expression changes across age groups for each cell type, we observed that scEssentials’ dysregulation only occurred in a small proportion of cell types (*median* 4/136) while *Pts, Tial1, Kras, Stub1* and *Lamp1* were the most actively changing scEssentials in the ageing group (Figure 6A). To understand whether ageing-related changes are more associated with certain cell types or tissues, we investigated the frequency of the scEssentials that showed significant dysregulation in ageing within each cell type. Strikingly, all cell types were found to have at least one scEssentials gene that significantly changed in age groups (Supplementary Figure 13C), with median scEssentials per cell type at 25 across three age groups. A barplot illustrated the top 10 cell types with the highest number of significant scEssentials genes associated (Figure 6B). Skeletal muscle satellite cells, T cells and epithelial cells were identified regardless of different tissues origins, which have shown profound function deterioration in ageing (48, 49). In addition, high numbers of differentially expressed scEssentials were found in closely interacted brain cell types such as neurons and oligodendrocytes. Studies have suggested that the oligodendroglia changes and their impacts on neurofunctional are highly associated with brain function declines in ageing (50).

## Discussion

Our work extends existing knowledge of essential genes to single-cell resolution at different molecular levels within the mouse and human organisms. Leveraging benchmarking datasets with more than 10 sequencing platforms and large-scale mouse and human transcription atlases, we identified robust scEssentials gene signatures across 60 cell types that can be used as a reference source in single-cell transcriptomic research. These genes are sought to be involved in basic cellular functions regardless of cell type. We confirmed this with a comprehensive study on the scEssentials at a single-gene, gene-pairs and multi-gene as well as multi-molecular level, improving the understanding of the gene essentiality in heterogeneous cellular functions.

We showed that scEssentials genes have high expression, high proportions of expression and non-cell-type-specificity in both human and mouse organisms. Because scEssentials genes characteristics are different to the cell-type-specific markers in scRNA-seq data, they can serve as a negative control for evaluation and comparison in marker identification tools. However, a distinguishing feature of scEssentials that separate them from housekeeping genes and/or SEGs, is that a level of gene expression variability is observed in scEssentials genes. Although such variation is significantly lower than in non-essential and non-housekeeping genes, it may reflect the transcriptional fluctuations under perturbations to maintain key functions specifically in human samples. Hence, scEssentials genes can be used for inferring cellular functions rather than defining stable and/or “control” gene expression in scRNA-seq data.

Our study unlocks the cell-state-specificity of scEssentials by investigating the gene-pair correlation across more than 60 cell types. Genes work together and regulate each other’s activity, thus, gene-gene correlation captures the underlying dynamics of genes’ regulation. Overall, a significantly higher number of correlated gene pairs and higher correlation in scEssentials genes compared to non-essential genes further support the critical co-regulation beyond a single gene level. Moreover, the consistently high coexpression patterns in the cell types in the specialised epithelial cells suggest the high regulatory constraints of the scEssentials genes on the differentiated cells. Our analysis further expands the knowledge that such tightly co-regulated and essential processes in specialised epithelial cells can be identified with different tissue origins, indicating scEssentials gene pairs are critical regulators to identify cell-type-specificity.

The consensus co-expression network constructed from various cell types has shown critical roles in pathways related to cell energy, cell cycle and cell maintenance. Each of the network communities is distinct and disjoint, demonstrating the unique roles of the selected scEssentials genes to maintain cellular functions. Incorporating consensus co-expression patterns into scRNA-seq clustering methods has already been developed to improve accuracy in identifying enriched pathways (51, 52). While these methods use different statistical approaches to find robust and denoised co-expression patterns across batches or biological conditions, it may be beneficial to include inherently stable co-expressed gene signatures from scEssentials genes to develop more accurate methods for clustering scRNA-seq data.

The essentiality score we developed showed strong associations with the degree of gene mutation and chromatin accessibility, demonstrating the robustness of the score at a multi-molecular level. The criticality that it illustrates allows for generating a flexible and robust quantification for any analytical applications in a data-dependent manner. Moreover, it provides evidence for the robust transcription process for the scEssentials with tight chromatin regulation.

In addition, our analysis demonstrates the high involvement of scEssentials genes in multiple pathways that are curated by two different pathway databases. Together with previous findings that essential genes are more likely to present as hub genes location in the protein-protein interactions (5), our result demonstrates crucial and key roles in the pathway networks location. It illustrates that scEssentials are the common regulatory genes to support complex biological processes. Furthermore, scEssentials can be used as one of the key sources when mitigating the highly overlapping genes among different pathways, which has been shown to improve the pathway overrepresentation analysis (53).

To further explore the robust gene expression of scEssentials, we investigated the scEssentials expression changes in the heterogeneous ageing process. We found that over 80% of the scEssentials genes do not significantly change their expression among the three age groups. This analysis demonstrates the robustness of scEssentials genes expression in the accumulated stress conditions. In contrast, the consistently differentially expressed scEssentials genes among different cell types have already been suggested to be dysregulated in ageing human skeletal muscle (e.g., *Pts*) or triggering upregulation of other ageing factors like p53 (e.g., *Tial1*) (54, 55). Therefore, such dysregulated scEssentials may be more likely to cause phenotypic changes in the cell types. Supporting our previous findings on the scEssentials gene expression that exist more variation in muscle satellite cells, our analysis found that these cell types had most of the differential expression scEssentials genes in ageing. In line with the stem cell decline which is one of the known hallmarks of ageing, we showed that stem cell extortion is a universal consequence in multiple stem-like cell types and constitutes multi-tissue and multi-organismal ageing (56).

Overall, our study demonstrates the significant role of scEssentials genes in maintaining cellular functions in multiple layers. Leveraging atlas-size scRNA-seq databases, we show robust, consistent, and high gene expression of scEssentials that can be used in different applications in analysing complex processes. Given the fundamental roles that scEssentials play, we demonstrate subtle changes in scEssential genes in ageing may cause significant consequences, especially in skeletal muscle satellite cells, T cells and epithelial cells of varying tissue origins.

