## Supplemental Info for "Assessing tissue-specific gene expression of essential genes from human and mouse"

**Supplementary Figures and Tables**

Supplementary Tables 1. Details of the benchmarking datasets used to measure the scEssential genes' expression detectability across sequencing methodology.

| **Organism** | **Number of cells** | **Sequencing method** | **Cell type** | **3' or full-length** | **Reference** |
|---|---|---|---|---|---|
| **Mouse** | 71 | CELseq2 | Embryonic stem cells | 3' | (1) |
| | 76 | Dropseq | | 3' | |
| | 65 | MARSeq | | 3' | |
| | 84 | SCRBseq | | 3' | |
| | 130 | Smartseq | | full-length | |
| | 157 | Smartseq2 | | full-length | |
| **Human** | 1439 | 10X Chromium (10X_LLU) | B lymphocytes | 3' | (2) |
| | 3296 | 10X Chromium (10X_NCI) | | 3' | |
| | 3273 | 10X Chromium (10X_NCI_M) | | 3' | |
| | 241 | Fluidigm C1 HT ( C1_FDA_HT) | | 3' | |
| | 66 | Fluidigm C1 ( C1_LLU) | | full-length | |
| | 596 | Takara Bio ICELL8 (ICELL8_PE) | | full-length | |
| | 600 | Takara Bio ICELL8 (ICELL8_SE) | | full-length | |

Supplementary Tables 2. The number of scEssentials and other protein-coding genes that had been classified based on gene mutation index. The *Chi-square test* was applied to determine the relationship ($p < 0.05$).

| | **scEssentials** | **Non-scEssentials** |
|---|---|---|
| **Low** | 111 | 316 |
| **Medium** | 4203 | 14121 |
| **High** | 112 | 695 |

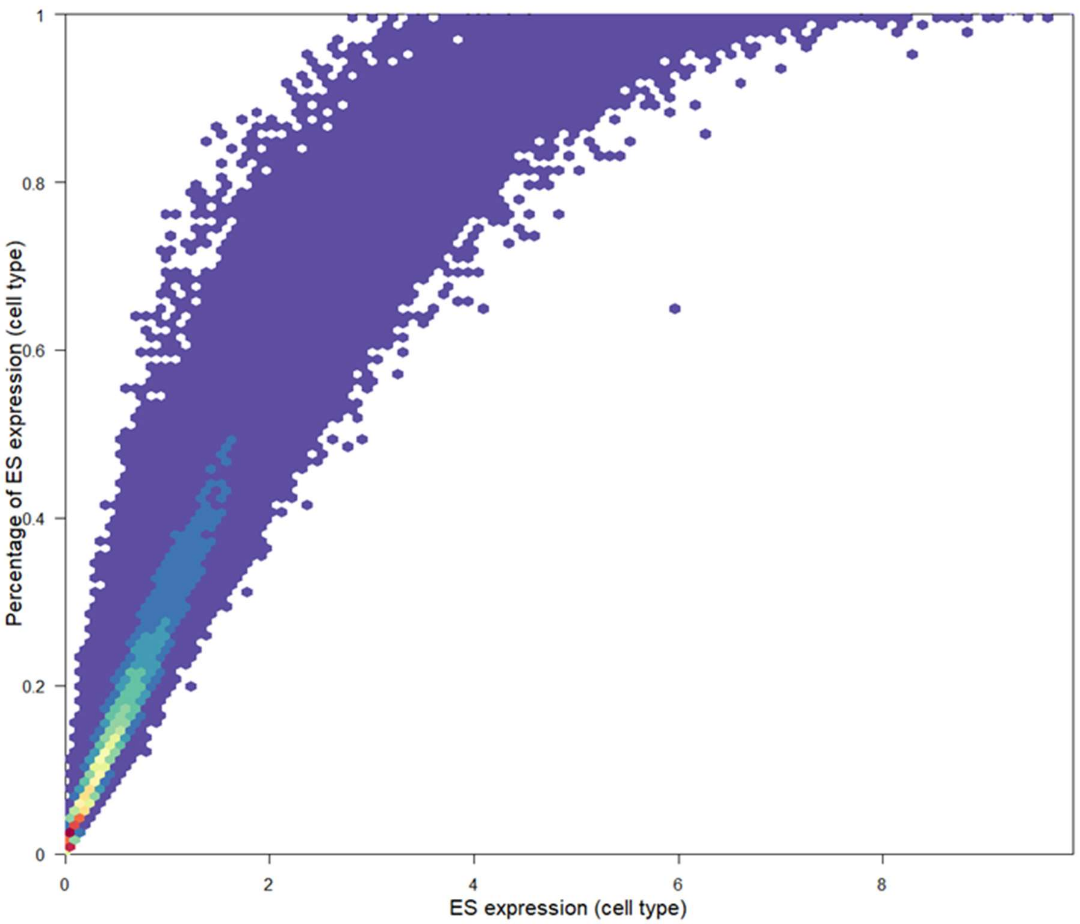


Supplementary Figure 1. Correlation of average expression and percentage of the cell’s expressed for scEssential genes. The high correlation illustrated the percentage of cell’s expression captured the high level of expression characteristics while providing a wider comparison range.

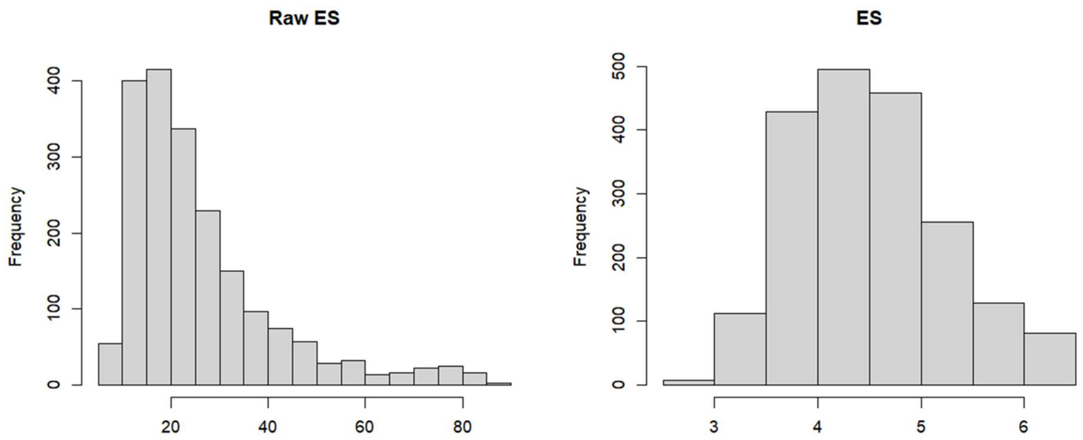


Supplementary Figure 2. The distribution of raw ES and ES values for scEssentials. On the left, raw ES distribution showed right-skewness and with substaintial value difference. To mitigate such skewness, logarithm2 transformation was applied on raw ES score.

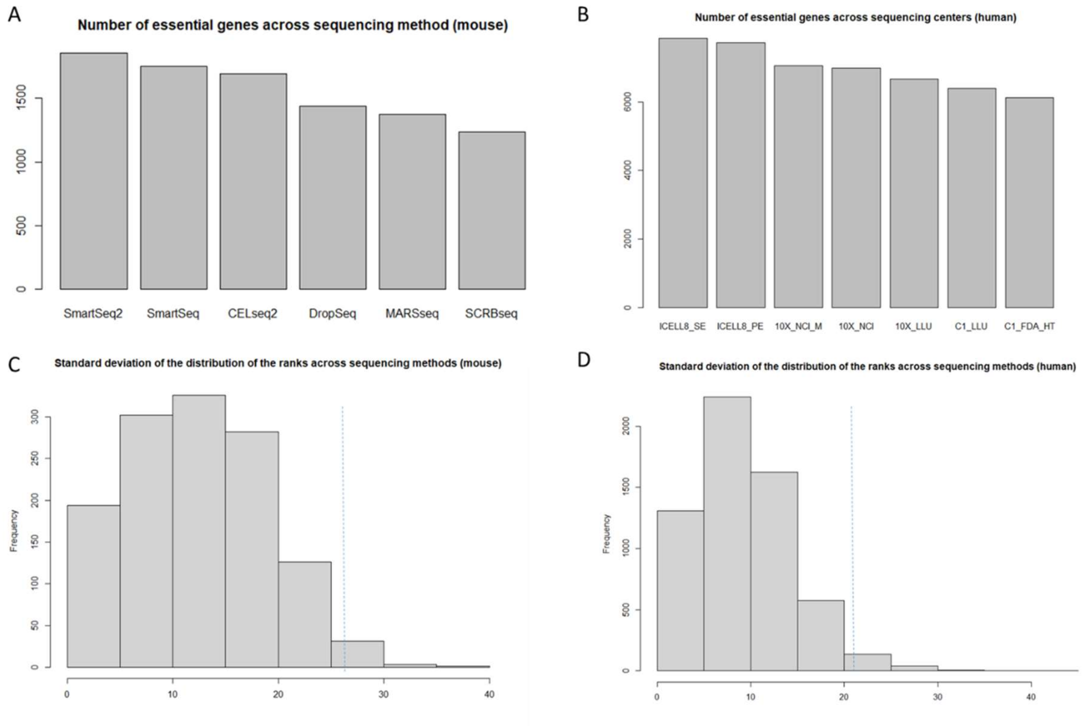


Supplementary Figure 3. Detectability and variations of essential genes across different sequencing platforms. The number of essential genes that had expression across A) six sequencing methods in mESCs (1) and B) seven sequencing methods (sites) in a matched control 'normal' B lymphocyte line from breast cancer patients. Figure C) and D) demonstrated the standard deviation (SD) of each essential gene's expression across sequencing for mESCs and human B cells. The blue dash lines represented 4 times the overall SD.

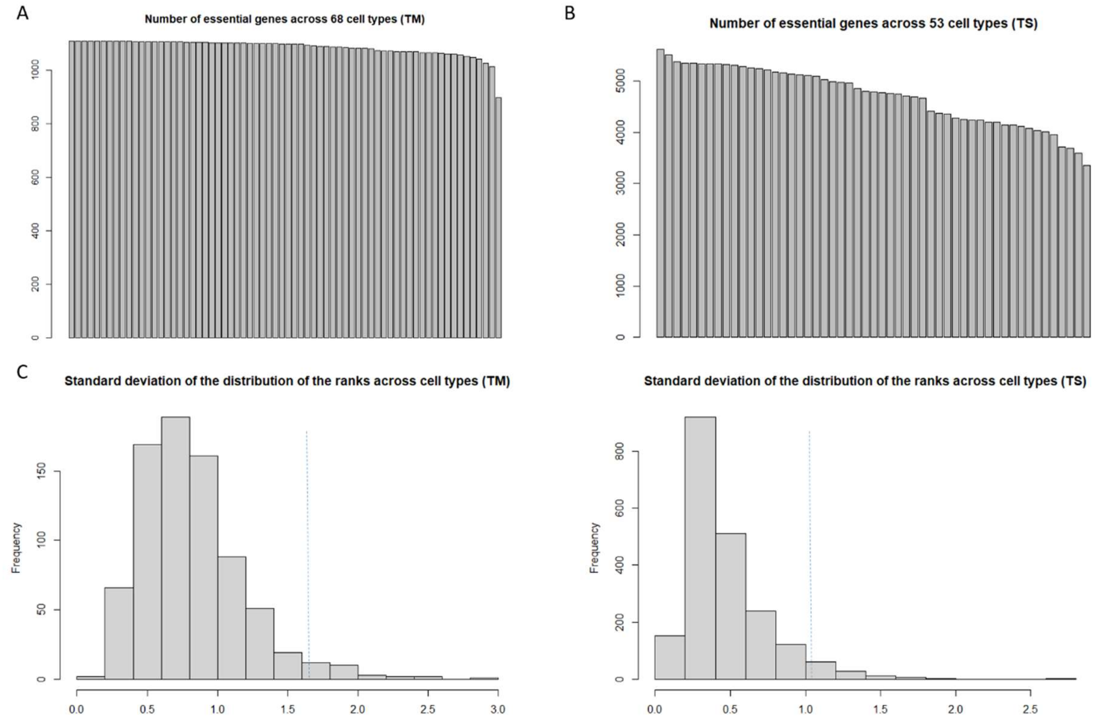


Supplementary Figure 4. Detectability and variations of essential genes in more than 60 unique cell types. The number of essential genes that had expression across A) 68 cell types in Tabula Muris (TM) and B) 53 cell types in Tabula Sapiens (TS) (3). Figure C and D demonstrated the standard deviation (SD) of each essential gene's expression across cell types in TM and TS. The blue dash lines represented 4 times the overall SD.

A

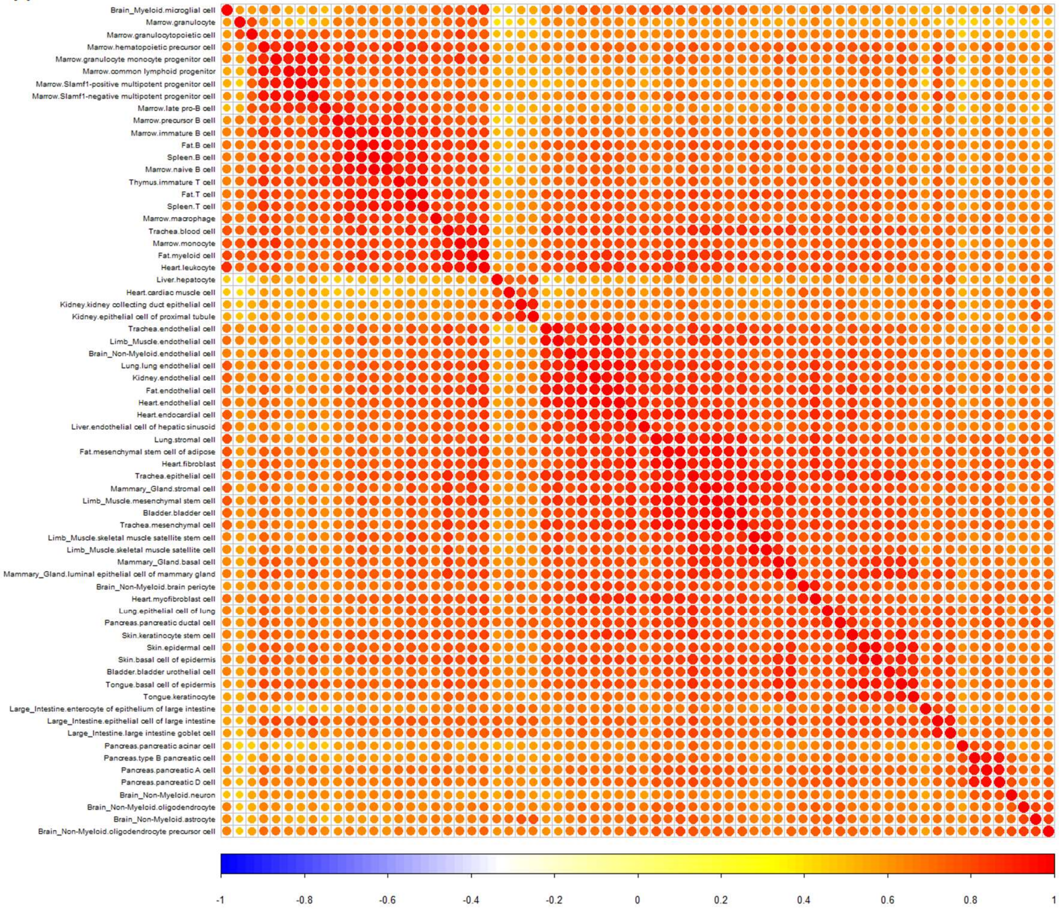

Brain_Myeloid.microglial cell
Marrow.granulocyte
Marrow.granulocytopoietic cell
Marrow.hematopoietic precursor cell
Marrow.granulocyte monocyte progenitor cell
Marrow.common lymphoid progenitor
Marrow.Slamf1-positive multipotent progenitor cell
Marrow.Slamf1-negative multipotent progenitor cell
Marrow.late pro-B cell
Marrow.precursor B cell
Marrow.immature B cell
Fat.B cell
Spleen.B cell
Marrow.naive B cell
Thymus.immature T cell
Fat.T cell
Spleen.T cell
Marrow.macrophage
Trachea.blood cell
Marrow.monocyte
Fat.myeloid cell
Heart.leukocyte
Liver.hepatocyte
Heart.cardiac muscle cell
Kidney.kidney collecting duct epithelial cell
Kidney.epithelial cell of proximal tubule
Trachea.endothelial cell
Limb_Muscle.endothelial cell
Brain_Non-Myeloid.endothelial cell
Lung.lung endothelial cell
Kidney.endothelial cell
Fat.endothelial cell
Heart.endothelial cell
Heart.endocardial cell
Liver.endothelial cell of hepatic sinusoid
Lung.stromal cell
Fat.mesenchymal stem cell of adipose
Heart.fibroblast
Trachea.epithelial cell
Mammary_Gland.stromal cell
Limb_Muscle.mesenchymal stem cell
Bladder.bladder cell
Trachea.mesenchymal cell
Limb_Muscle.skeletal muscle satellite stem cell
Limb_Muscle.skeletal muscle satellite cell
Mammary_Gland.basal cell
Mammary_Gland.luminal epithelial cell of mammary gland
Brain_Non-Myeloid.brain pericyte
Heart.myofibroblast cell
Lung.epithelial cell of lung
Pancreas.pancreatic ductal cell
Skin.keratinocyte stem cell
Skin.epidermal cell
Skin.basal cell of epidermis
Bladder.bladder urothelial cell
Tongue.basal cell of epidermis
Tongue.keratinocyte
Large_Intestine.enterocyte of epithelium of large intestine
Large_Intestine.epithelial cell of large intestine
Large_Intestine.large intestine goblet cell
Pancreas.pancreatic acinar cell
Pancreas.type B pancreatic cell
Pancreas.pancreatic A cell
Pancreas.pancreatic D cell
Brain_Non-Myeloid.neuron
Brain_Non-Myeloid.oligodendrocyte
Brain_Non-Myeloid.astrocyte
Brain_Non-Myeloid.oligodendrocyte precursor cell
-1
-0.8
-0.6
-0.4
-0.2
0
0.2
0.4
0.6
0.8
1

B

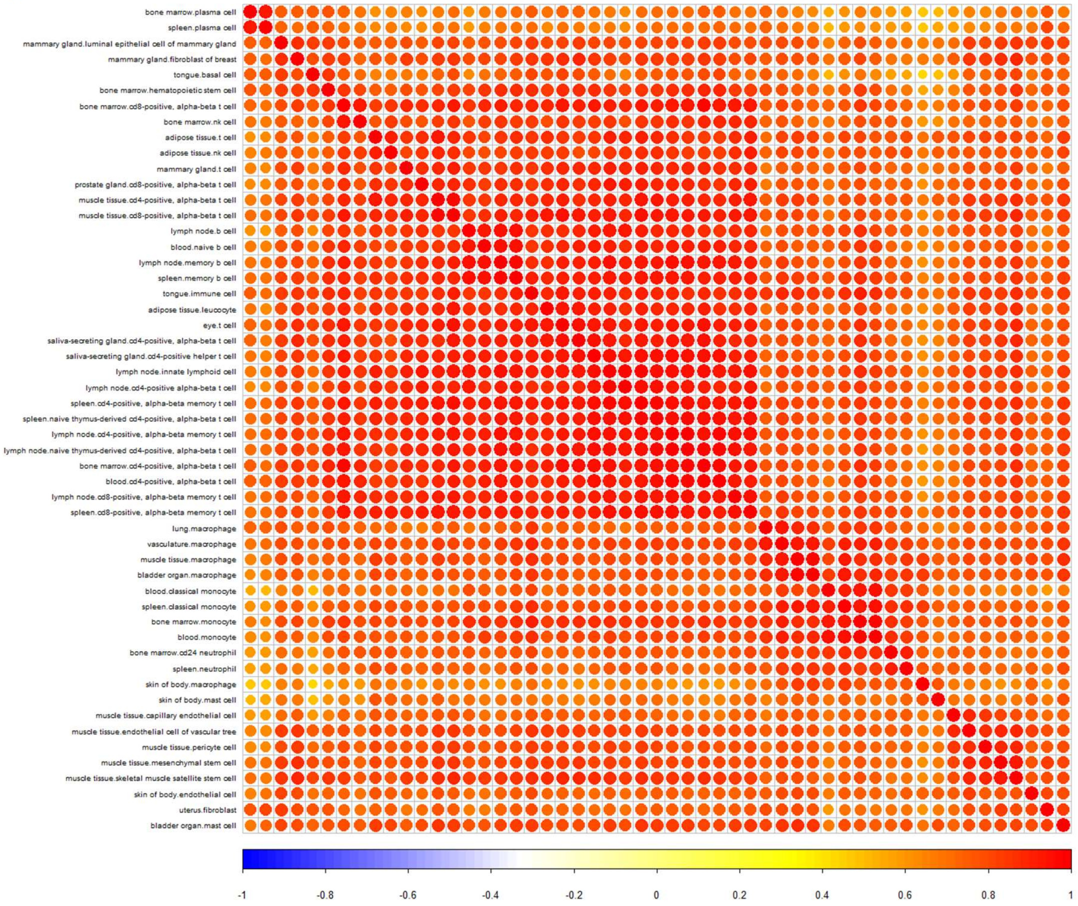


Supplementary Figure 5. Correlation heatmap across cell types based on scEssential genes. Pearson correlation was applied to measure the similarities among cell types. By performing the correlation with expression values, the heatmaps showed non-cell-type-specific patterns with an average higher correlation A) with the TM dataset; B) with the TS dataset. But the correlation reduced with clear cell-type-specificity when the scEssentials expression ranking replaced the expression value.

A

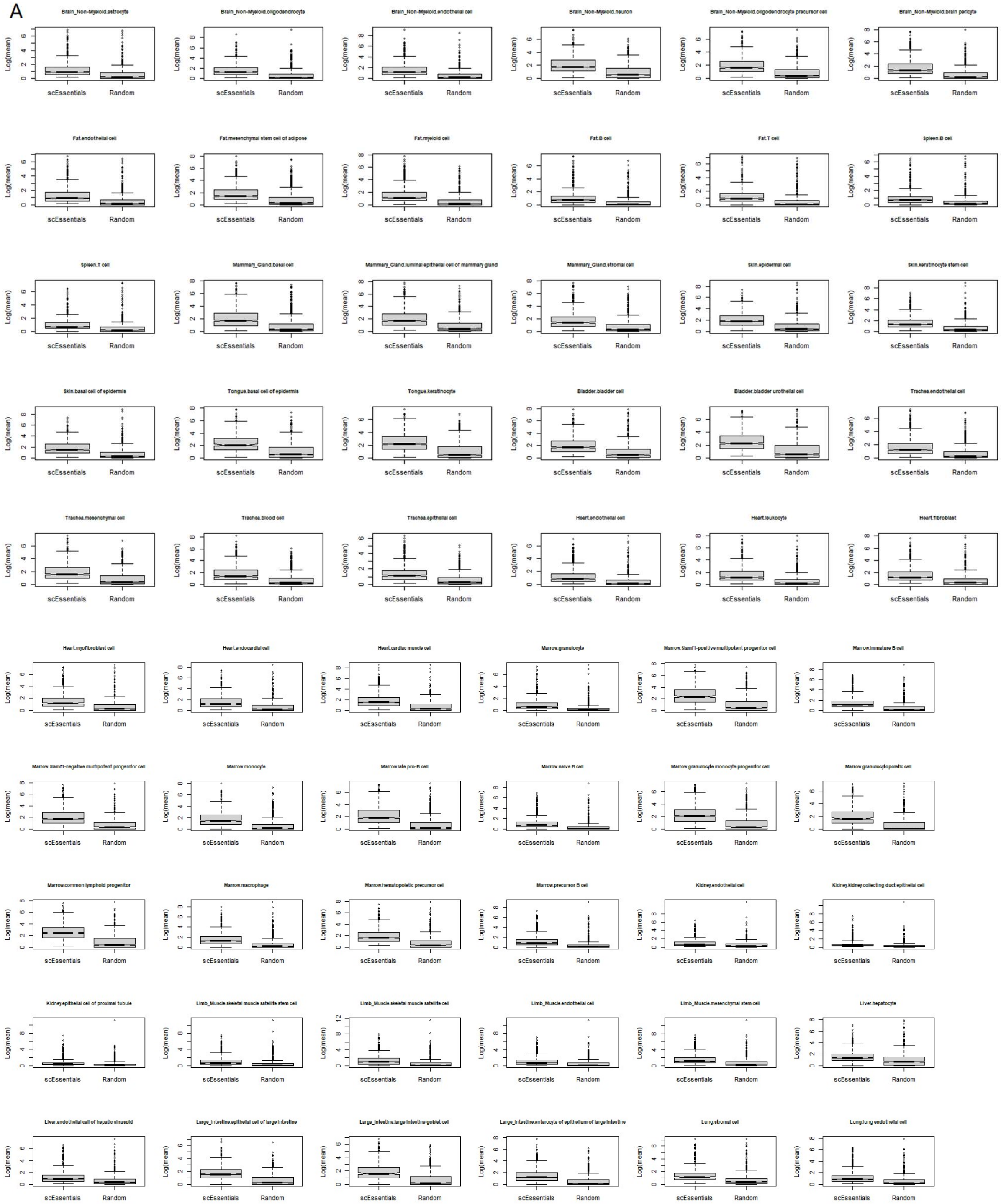
Brain_Non-Myeloid.astrocyte
Brain_Non-Myeloid.oligodendrocyte
Brain_Non-Myeloid.endothelial cell
Brain_Non-Myeloid.neuron
Brain_Non-Myeloid.oligodendrocyte precursor cell
Brain_Non-Myeloid.brain pericyte
Fat.endothelial cell
Fat.mesenchymal stem cell of adipose
Fat.myeloid cell
Fat.B cell
Fat.T cell
Spleen.B cell
Spleen.T cell
Mammary_Gland.basal cell
Mammary_Gland.luminal epithelial cell of mammary gland
Mammary_Gland.stromal cell
Skin.epidermal cell
Skin.keratinocyte stem cell
Skin.basal cell of epidermis
Tongue.basal cell of epidermis
Tongue.keratinocyte
Bladder.bladder cell
Bladder.bladder urothelial cell
Trachea.endothelial cell
Trachea.mesenchymal cell
Trachea.blood cell
Trachea.epithelial cell
Heart.endothelial cell
Heart.leukocyte
Heart.fibroblast
Heart.myofibroblast cell
Heart.endocardial cell
Heart.cardiac muscle cell
Marrow.granulocyte
Marrow.Slamf1-positive multipotent progenitor cell
Marrow.immature B cell
Marrow.Slamf1-negative multipotent progenitor cell
Marrow.monocyte
Marrow.late pro-B cell
Marrow.naive B cell
Marrow.granulocyte monocyte progenitor cell
Marrow.granulocytopoietic cell
Marrow.common lymphoid progenitor
Marrow.macrophage
Marrow.hematopoietic precursor cell
Marrow.precursor B cell
Kidney.endothelial cell
Kidney.kidney collecting duct epithelial cell
Kidney.epithelial cell of proximal tubule
Limb_Muscle.skeletal muscle satellite stem cell
Limb_Muscle.skeletal muscle satellite cell
Limb_Muscle.endothelial cell
Limb_Muscle.mesenchymal stem cell
Liver.hepatocyte
Liver.endothelial cell of hepatic sinusoid
Large_Intestine.epithelial cell of large intestine
Large_Intestine.large intestine goblet cell
Large_Intestine.enterocyte of epithelium of large intestine
Lung.stromal cell
Lung.lung endothelial cell
Log(mean)
scEssentials
Random

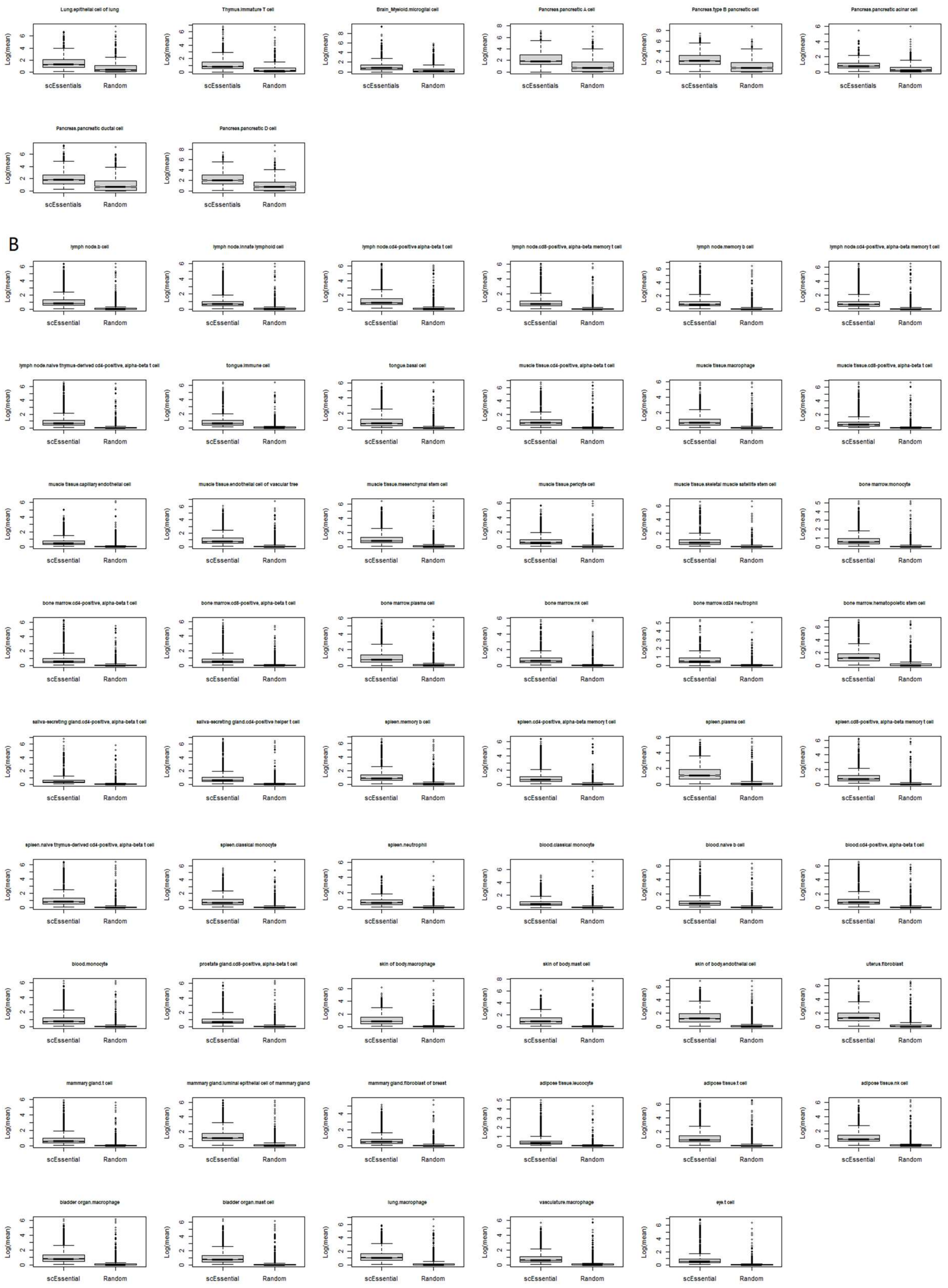


Supplementary Figure 6. The significantly high average expression in scEssentials as compared to a random gene list. The Wilcoxon ranked test was applied for all cell types to compare the log of the mean expression difference in A) TM and B) TS. All cell types under various comparisons showed a significant increase in scEssentials with respect to the random gene list (Wilcoxon rank test, $p<0.05$).

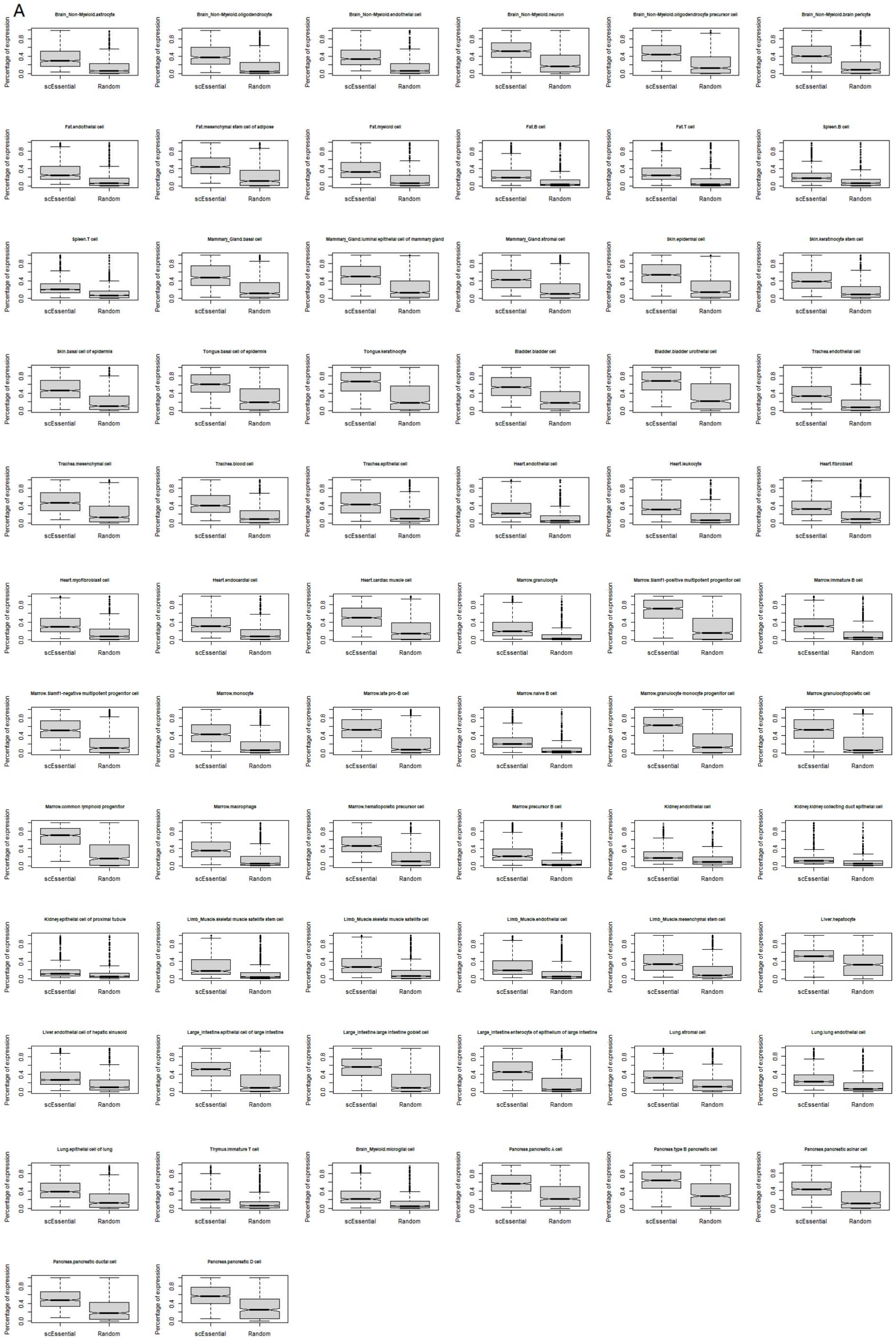
A
Brain_Non-Myeloid.astrocyte
Brain_Non-Myeloid.oligodendrocyte
Brain_Non-Myeloid.endothelial cell
Brain_Non-Myeloid.neuron
Brain_Non-Myeloid.oligodendrocyte precursor cell
Brain_Non-Myeloid.brain pericyte
Fat.endothelial cell
Fat.mesenchymal stem cell of adipose
Fat.myeloid cell
Fat.B cell
Fat.T cell
Spleen.B cell
Spleen.T cell
Mammary_Gland.basal cell
Mammary_Gland.luminal epithelial cell of mammary gland
Mammary_Gland.stromal cell
Skin.epidermal cell
Skin.keratinocyte stem cell
Skin.basal cell of epidermis
Tongue.basal cell of epidermis
Tongue.keratinocyte
Bladder.bladder cell
Bladder.bladder urothelial cell
Trachea.endothelial cell
Trachea.mesenchymal cell
Trachea.blood cell
Trachea.epithelial cell
Heart.endothelial cell
Heart.leukocyte
Heart.fibroblast
Heart.myofibroblast cell
Heart.endocardial cell
Heart.cardiac muscle cell
Marrow.granulocyte
Marrow.Slamf1-positive multipotent progenitor cell
Marrow.immature B cell
Marrow.Slamf1-negative multipotent progenitor cell
Marrow.monocyte
Marrow.late pro-B cell
Marrow.naive B cell
Marrow.granulocyte monocyte progenitor cell
Marrow.granulocytopoietic cell
Marrow.common lymphoid progenitor
Marrow.macrophage
Marrow.hematopoietic precursor cell
Marrow.precursor B cell
Kidney.endothelial cell
Kidney.kidney collecting duct epithelial cell
Kidney.epithelial cell of proximal tubule
Limb_Muscle.skeletal muscle satellite stem cell
Limb_Muscle.skeletal muscle satellite cell
Limb_Muscle.endothelial cell
Limb_Muscle.mesenchymal stem cell
Liver.hepatocyte
Liver.endothelial cell of hepatic sinusoid
Large_Intestine.epithelial cell of large intestine
Large_Intestine.large intestine goblet cell
Large_Intestine.enterocyte of epithelium of large intestine
Lung.stromal cell
Lung.lung endothelial cell
Lung.epithelial cell of lung
Thymus.immature T cell
Brain_Myeloid.microglial cell
Pancreas.pancreatic A cell
Pancreas.type B pancreatic cell
Pancreas.pancreatic acinar cell
Pancreas.pancreatic ductal cell
Pancreas.pancreatic D cell
Percentage of expression
0.0
0.4
0.8
scEssential
Random

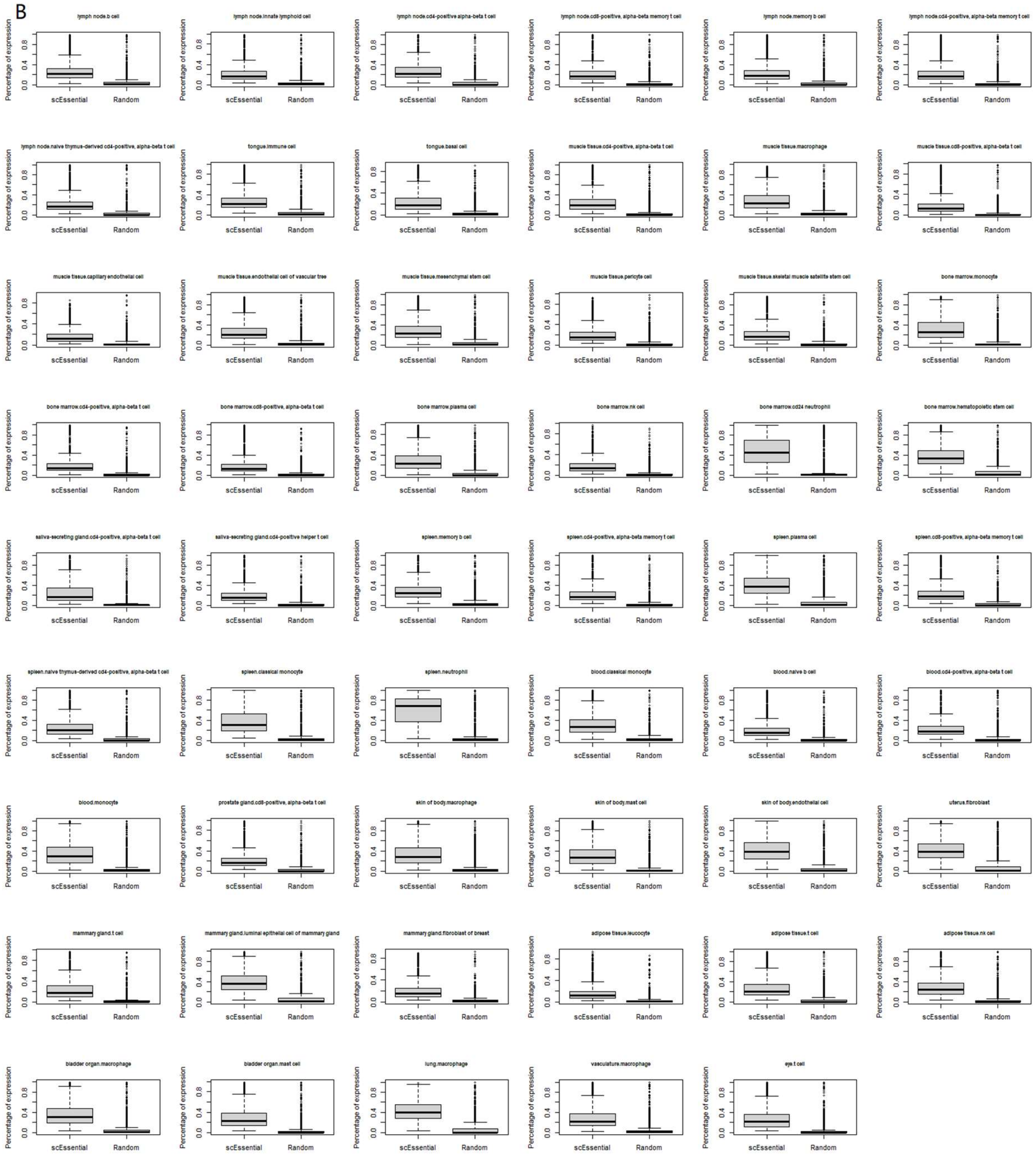


Supplementary Figure 7. The significantly high percentage of cells expressed in scEssentials as compared to a random gene list. The Wilcoxon ranked test was applied for all cell types to compare the percentage of expression difference in A) TM and B) TS. All cell types under various comparisons showed a significant increase in scEssentials with respect to the random gene list (Wilcoxon rank test, p<0.05).

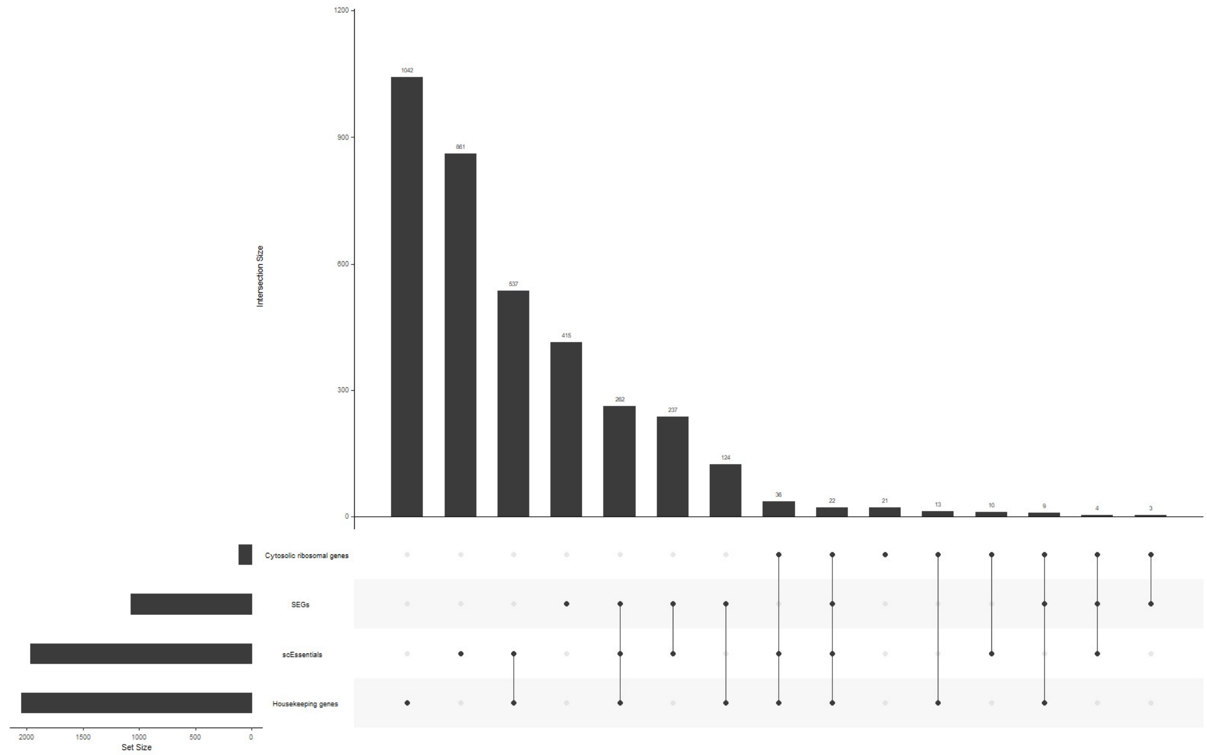


Supplementary Figure 8. The upset plot demonstrated the overlap between scEssentials and other gene lists with high stability and importance, including housekeeping genes, ribosomal genes and stably expressed genes (SEGs). Housekeeping genes were retrieved from the housekeeping genes database website (4). Cytosolic ribosomal genes were selected from *Deeke et al.,* (5) and stably expressed genes were retrieved from *Lin et al*., (6).

A

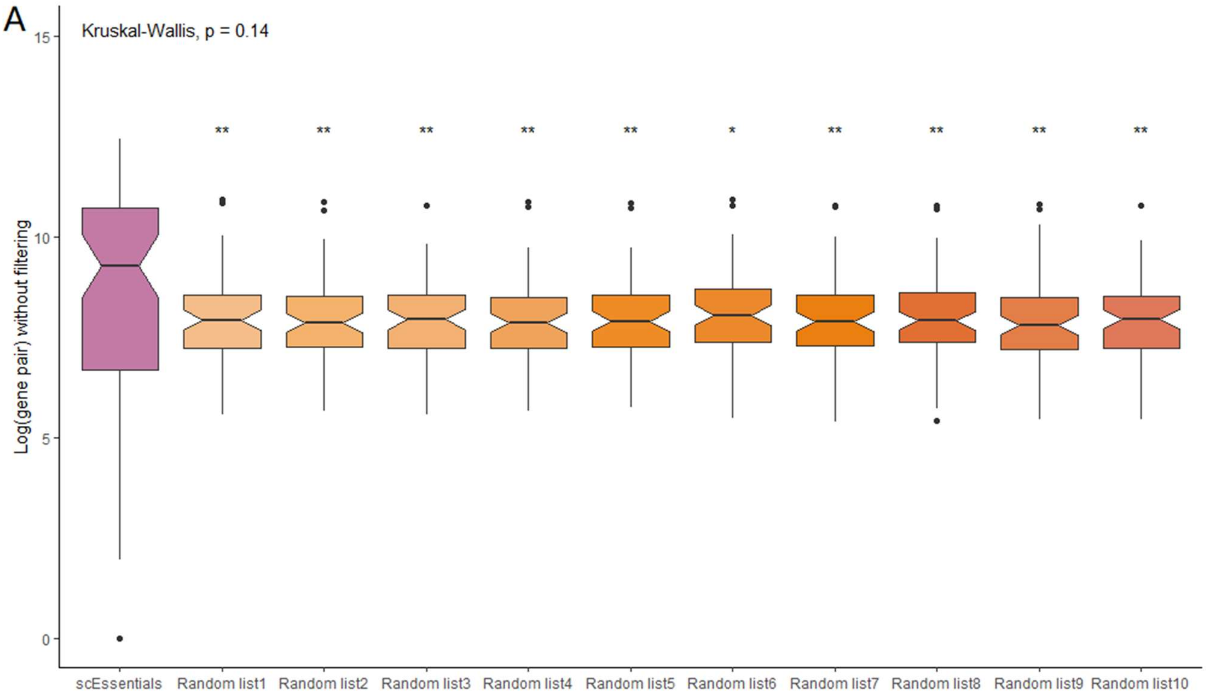

Kruskal-Wallis, p = 0.14
Log(gene pair) without filtering
15
10
5
0
scEssentials
Random list1
Random list2
Random list3
Random list4
Random list5
Random list6
Random list7
Random list8
Random list9
Random list10


B

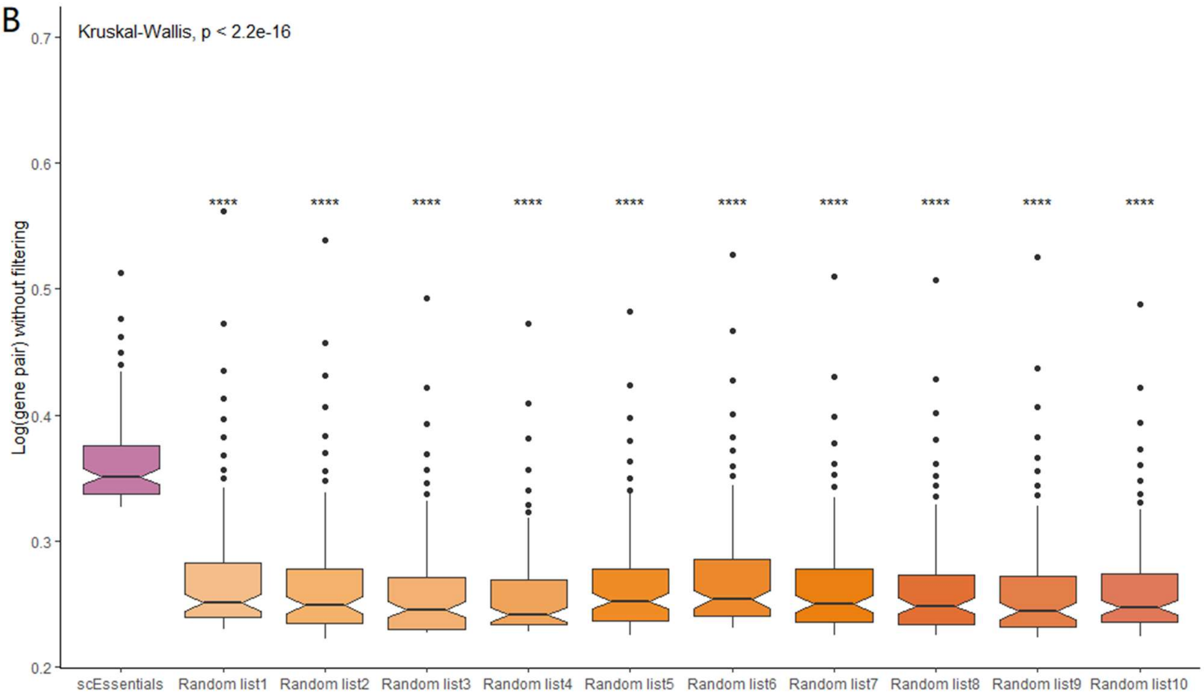

Kruskal-Wallis, p < 2.2e-16
Log(gene pair) without filtering
0.7
0.6
0.5
0.4
0.3
0.2
scEssentials
Random list1
Random list2
Random list3
Random list4
Random list5
Random list6
Random list7
Random list8
Random list9
Random list10

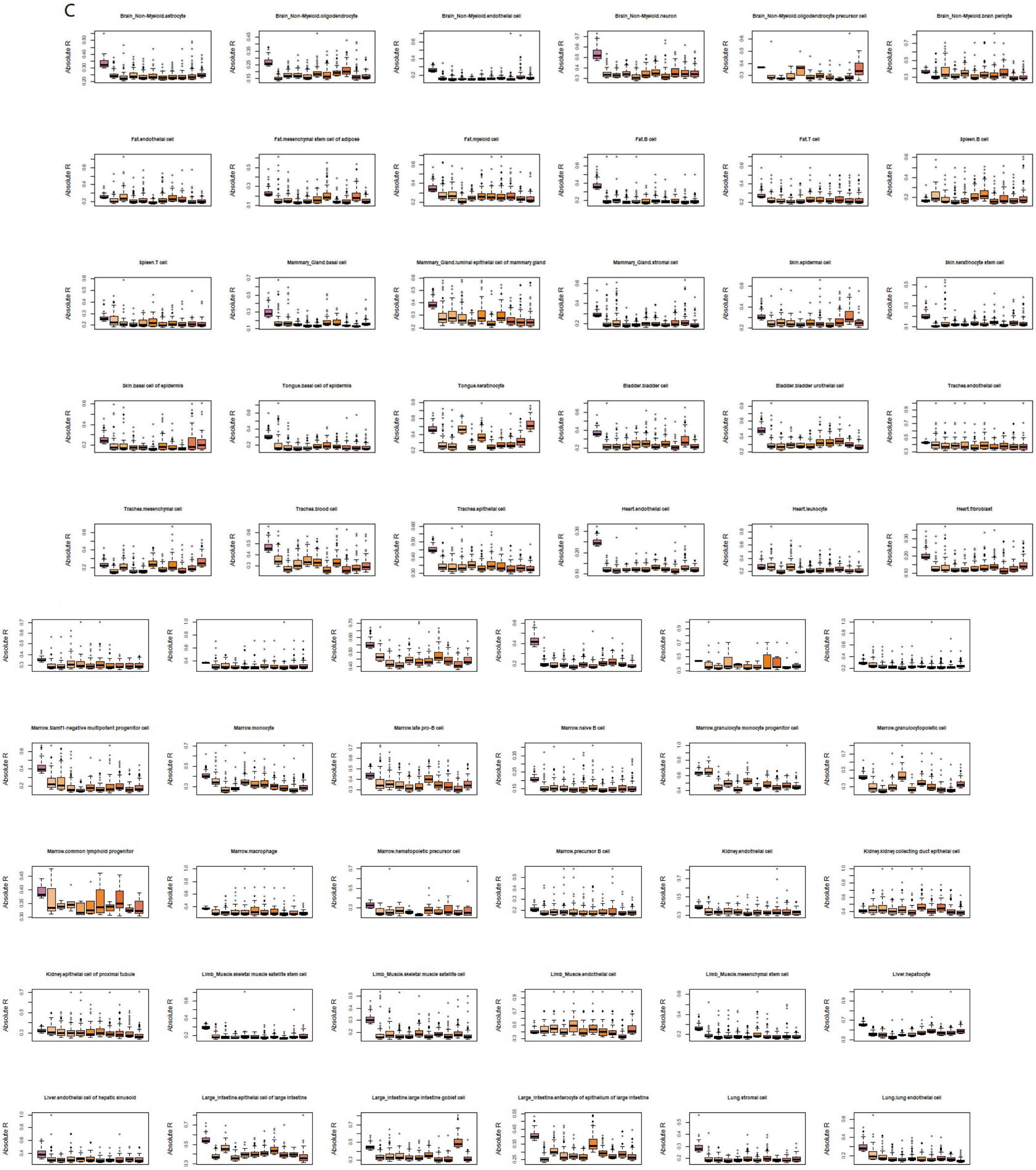
C
Brain_Non-Myeloid.astrocyte
Brain_Non-Myeloid.oligodendrocyte
Brain_Non-Myeloid.endothelial cell
Brain_Non-Myeloid.neuron
Brain_Non-Myeloid.oligodendrocyte precursor cell
Brain_Non-Myeloid.brain pericyte
Fat.endothelial cell
Fat.mesenchymal stem cell of adipose
Fat.myeloid cell
Fat.B cell
Fat.T cell
Spleen.B cell
Spleen.T cell
Mammary_Gland.basal cell
Mammary_Gland.luminal epithelial cell of mammary gland
Mammary_Gland.stromal cell
Skin.epidermal cell
Skin.keratinocyte stem cell
Skin.basal cell of epidermis
Tongue.basal cell of epidermis
Tongue.keratinocyte
Bladder.bladder cell
Bladder.bladder urothelial cell
Trachea.endothelial cell
Trachea.mesenchymal cell
Trachea.blood cell
Trachea.epithelial cell
Heart.endothelial cell
Heart.leukocyte
Heart.fibroblast
Marrow.Slamf1-negative multipotent progenitor cell
Marrow.monocyte
Marrow.late pro-B cell
Marrow.naive B cell
Marrow.granulocyte monocyte progenitor cell
Marrow.granulocytopoietic cell
Marrow.common lymphoid progenitor
Marrow.macrophage
Marrow.hematopoietic precursor cell
Marrow.precursor B cell
Kidney.endothelial cell
Kidney.kidney collecting duct epithelial cell
Kidney.epithelial cell of proximal tubule
Limb_Muscle.skeletal muscle satellite stem cell
Limb_Muscle.skeletal muscle satellite cell
Limb_Muscle.endothelial cell
Limb_Muscle.mesenchymal stem cell
Liver.hepatocyte
Liver.endothelial cell of hepatic sinusoid
Large_Intestine.epithelial cell of large intestine
Large_Intestine.large intestine goblet cell
Large_Intestine.enterocyte of epithelium of large intestine
Lung.stromal cell
Lung.lung endothelial cell
Absolute R

Supplementary Figure 9. Coexpression level of scEssentials gene compared to 10 randomly sampled gene lists. A) The number of significantly correlated gene pairs for scEssentials genes as compared to other 10 random gene lists. Boxplots for all the genes in each gene list without filtering. B) The absolute correlation of coefficient for the top 100 most significantly coexpressed gene pairs for scEssentials as compared to the other 10 random gene lists. Boxplots for all the genes in each gene list after filtering. *Kruskal-Wallis* test was applied to determine the significance (* $p<0.05$; ** $p<0.01$; *** $p<0.001$. **** $p<0.0001$). C) The absolute correlation of coefficient for the top 100 most significantly coexpressed gene pairs in 68 cell types. Notably, only Slamf1-positive multipotent progenitor cell and common lymphoid progenitor cells did not show significantly higher correlation efficient in scEssentials with respect to other randomly sampled gene lists.

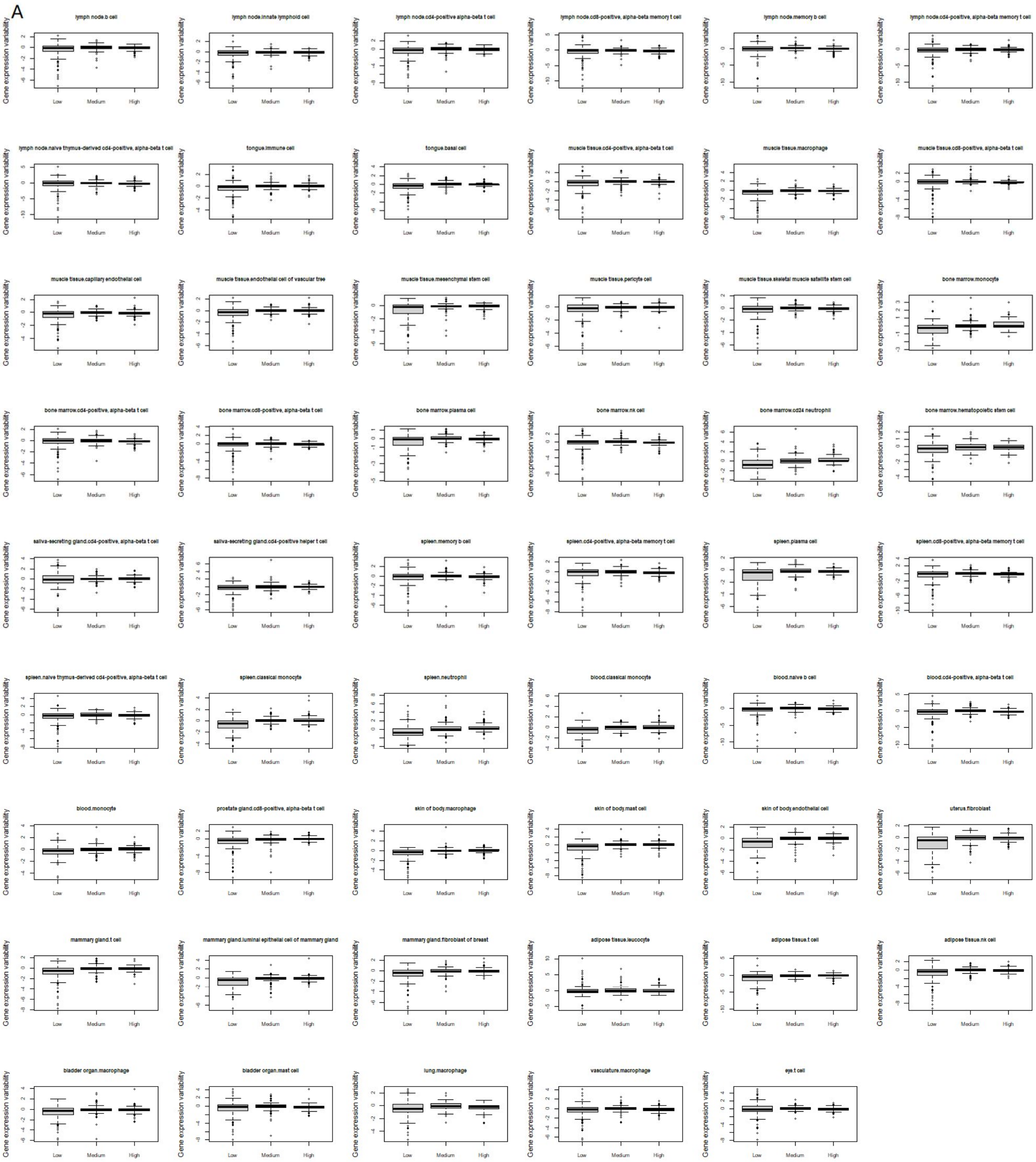
A
lymph node.b cell
lymph node.innate lymphoid cell
lymph node.cd4-positive alpha-beta t cell
lymph node.cd8-positive, alpha-beta memory t cell
lymph node.memory b cell
lymph node.cd4-positive, alpha-beta memory t cell
lymph node.naive thymus-derived cd4-positive, alpha-beta t cell
tongue.immune cell
tongue.basal cell
muscle tissue.cd4-positive, alpha-beta t cell
muscle tissue.macrophage
muscle tissue.cd8-positive, alpha-beta t cell
muscle tissue.capillary endothelial cell
muscle tissue.endothelial cell of vascular tree
muscle tissue.mesenchymal stem cell
muscle tissue.pericyte cell
muscle tissue.skeletal muscle satellite stem cell
bone marrow.monocyte
bone marrow.cd4-positive, alpha-beta t cell
bone marrow.cd8-positive, alpha-beta t cell
bone marrow.plasma cell
bone marrow.nk cell
bone marrow.cd24 neutrophil
bone marrow.hematopoietic stem cell
saliva-secreting gland.cd4-positive, alpha-beta t cell
saliva-secreting gland.cd4-positive helper t cell
spleen.memory b cell
spleen.cd4-positive, alpha-beta memory t cell
spleen.plasma cell
spleen.cd8-positive, alpha-beta memory t cell
spleen.naive thymus-derived cd4-positive, alpha-beta t cell
spleen.classical monocyte
spleen.neutrophil
blood.classical monocyte
blood.naive b cell
blood.cd4-positive, alpha-beta t cell
blood.monocyte
prostate gland.cd8-positive, alpha-beta t cell
skin of body.macrophage
skin of body.mast cell
skin of body.endothelial cell
uterus.fibroblast
mammary gland.t cell
mammary gland.luminal epithelial cell of mammary gland
mammary gland.fibroblast of breast
adipose tissue.leucocyte
adipose tissue.t cell
adipose tissue.nk cell
bladder organ.macrophage
bladder organ.mast cell
lung.macrophage
vasculature.macrophage
eye.t cell
Gene expression variability
Low
Medium
High

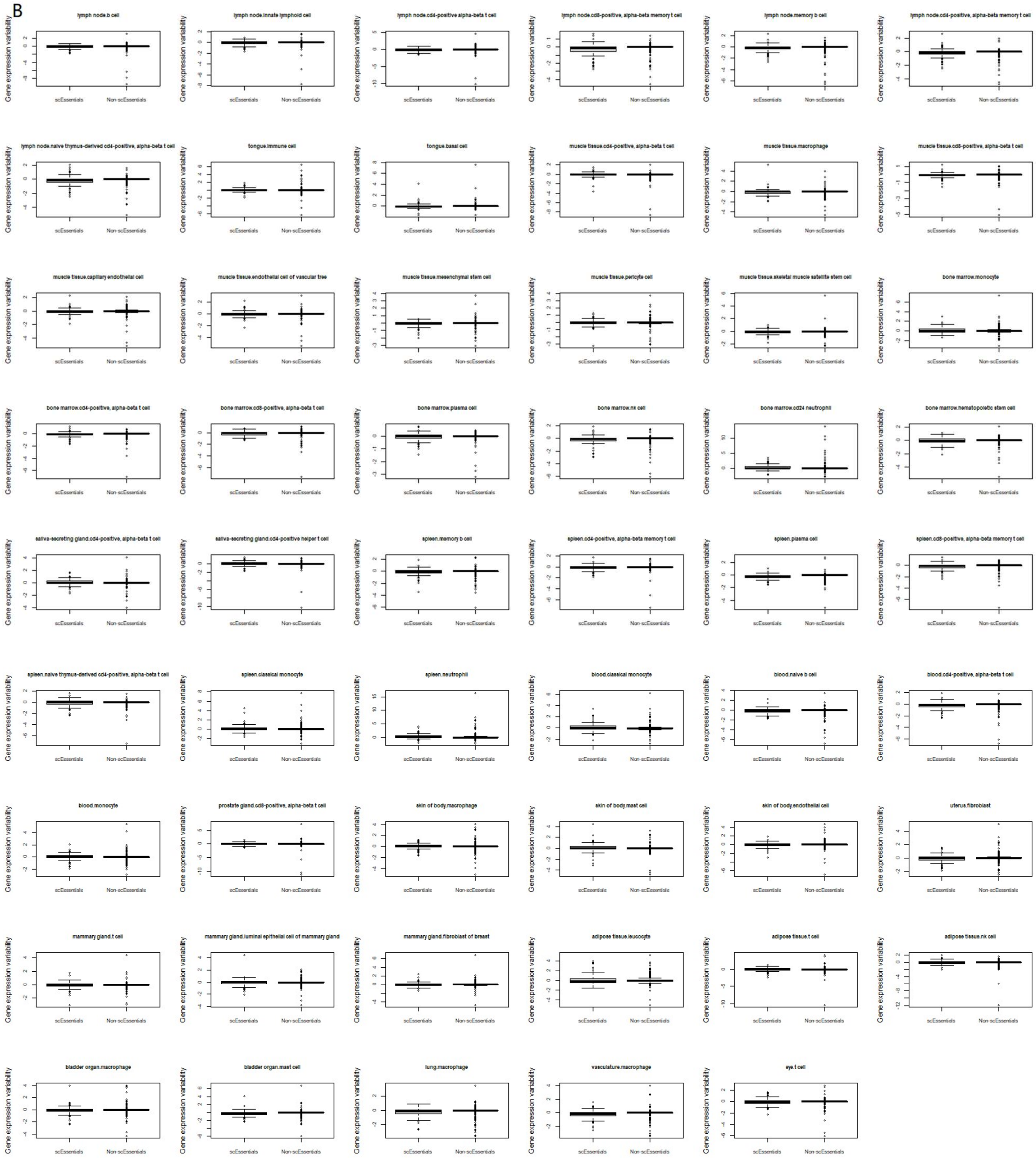
B
lymph node.b cell
lymph node.innate lymphoid cell
lymph node.cd4-positive alpha-beta t cell
lymph node.cd8-positive, alpha-beta memory t cell
lymph node.memory b cell
lymph node.cd4-positive, alpha-beta memory t cell
lymph node.naive thymus-derived cd4-positive, alpha-beta t cell
tongue.immune cell
tongue.basal cell
muscle tissue.cd4-positive, alpha-beta t cell
muscle tissue.macrophage
muscle tissue.cd8-positive, alpha-beta t cell
muscle tissue.capillary endothelial cell
muscle tissue.endothelial cell of vascular tree
muscle tissue.mesenchymal stem cell
muscle tissue.pericyte cell
muscle tissue.skeletal muscle satellite stem cell
bone marrow.monocyte
bone marrow.cd4-positive, alpha-beta t cell
bone marrow.cd8-positive, alpha-beta t cell
bone marrow.plasma cell
bone marrow.nk cell
bone marrow.cd24 neutrophil
bone marrow.hematopoietic stem cell
saliva-secreting gland.cd4-positive, alpha-beta t cell
saliva-secreting gland.cd4-positive helper t cell
spleen.memory b cell
spleen.cd4-positive, alpha-beta memory t cell
spleen.plasma cell
spleen.cd8-positive, alpha-beta memory t cell
spleen.naive thymus-derived cd4-positive, alpha-beta t cell
spleen.classical monocyte
spleen.neutrophil
blood.classical monocyte
blood.naive b cell
blood.cd4-positive, alpha-beta t cell
blood.monocyte
prostate gland.cd8-positive, alpha-beta t cell
skin of body.macrophage
skin of body.mast cell
skin of body.endothelial cell
uterus.fibroblast
mammary gland.t cell
mammary gland.luminal epithelial cell of mammary gland
mammary gland.fibroblast of breast
adipose tissue.leucocyte
adipose tissue.t cell
adipose tissue.nk cell
bladder organ.macrophage
bladder organ.mast cell
lung.macrophage
vasculature.macrophage
eye.t cell
Gene expression variability
scEssentials
Non-scEssentials

C

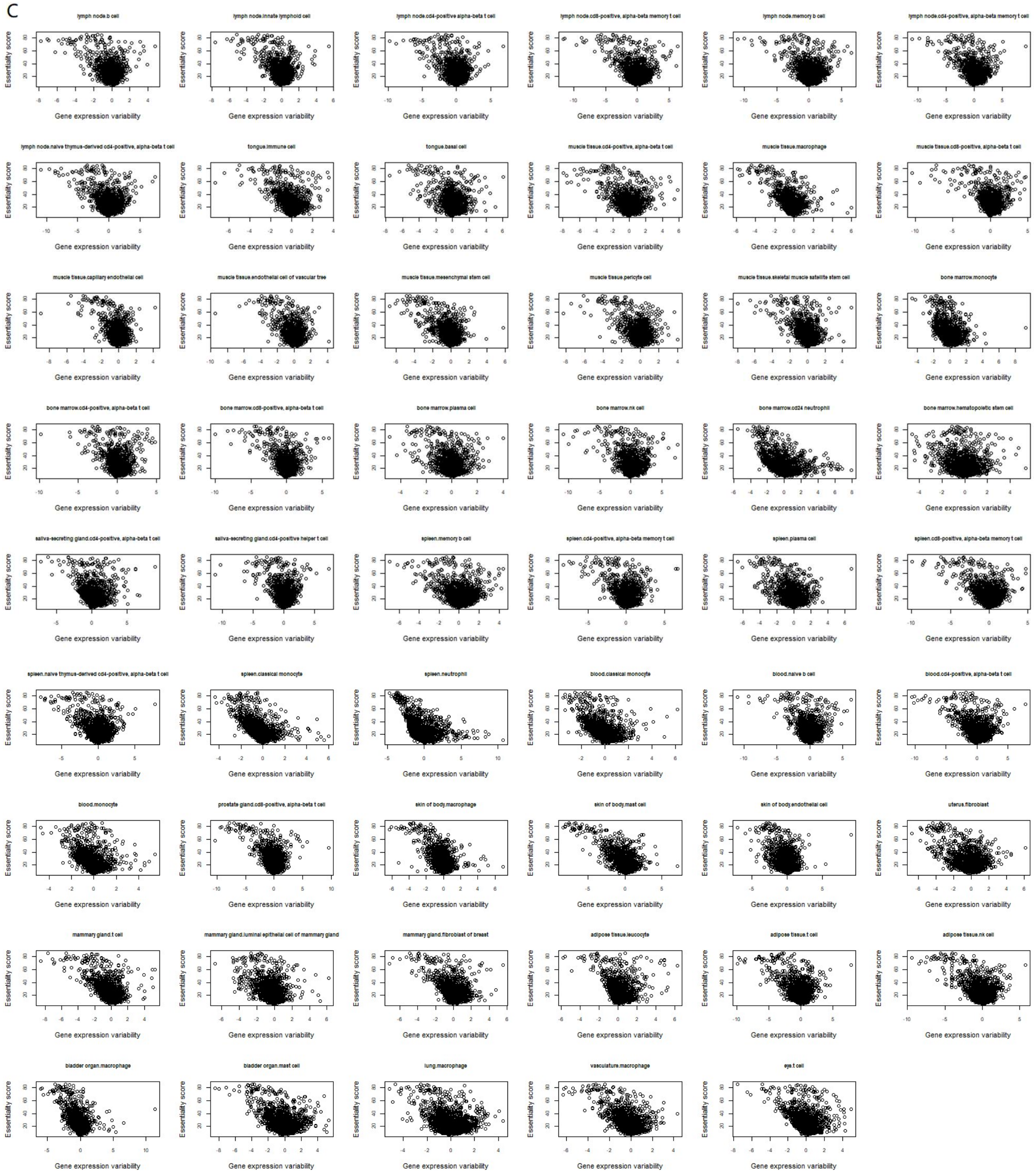


Supplementary Figure 10. Relationship of scEssential genes with different gene damage index (7). A) Boxplot for gene expression variability for each cell type under low, medium and high risks in each cell type. B) Boxplot for gene expression variability for each cell type between scEssentials and non-scEssentials under high risks. C) Correlation between ES and scEssentials expression variability across all cell types. The correlation coefficients were measured by Spearman correlation and were significantly correlated. The median correlation coefficient across all cell types was -0.4.

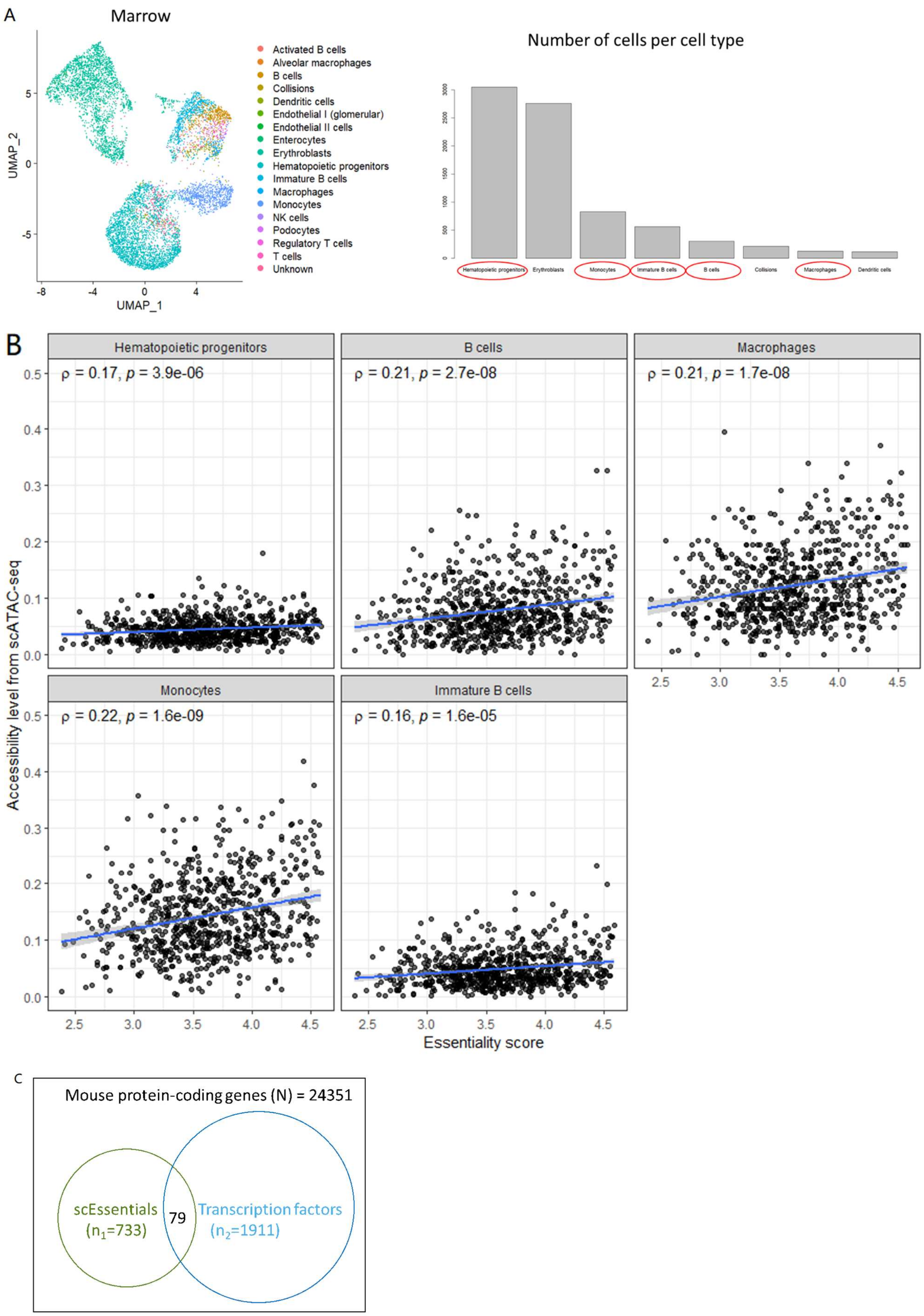
A
Marrow
Activated B cells
Alveolar macrophages
B cells
Collisions
Dendritic cells
Endothelial I (glomerular)
Endothelial II cells
Enterocytes
Erythroblasts
Hematopoietic progenitors
Immature B cells
Macrophages
Monocytes
NK cells
Podocytes
Regulatory T cells
T cells
Unknown
UMAP_1
UMAP_2
Number of cells per cell type
B
Hematopoietic progenitors
ρ = 0.17, p = 3.9e-06
B cells
ρ = 0.21, p = 2.7e-08
Macrophages
ρ = 0.21, p = 1.7e-08
Monocytes
ρ = 0.22, p = 1.6e-09
Immature B cells
ρ = 0.16, p = 1.6e-05
Accessibility level from scATAC-seq
Essentiality score
C
Mouse protein-coding genes (N) = 24351
scEssentials
(n1=733)
79
Transcription factors
(n2=1911)

Supplementary Figure 11. Chromatin accessibility inferred from scATAC-seq mouse atlas (12). A) UMAP demonstrated the clustering of the cell types in bone marrow tissue based on scATAC-seq data, and the cell types with more than 100 cells were plotted as barplot where the cell type that overlapped with TM data was circled. B) Dotplot demonstrated the correlation between the essentiality score and the accessibility level for each cell type and on average. Spearman correlation was applied to measure the correlation. Blue line represented the linear regression between two variables. C) Illustration of the parameters used for the hypergeometric test.

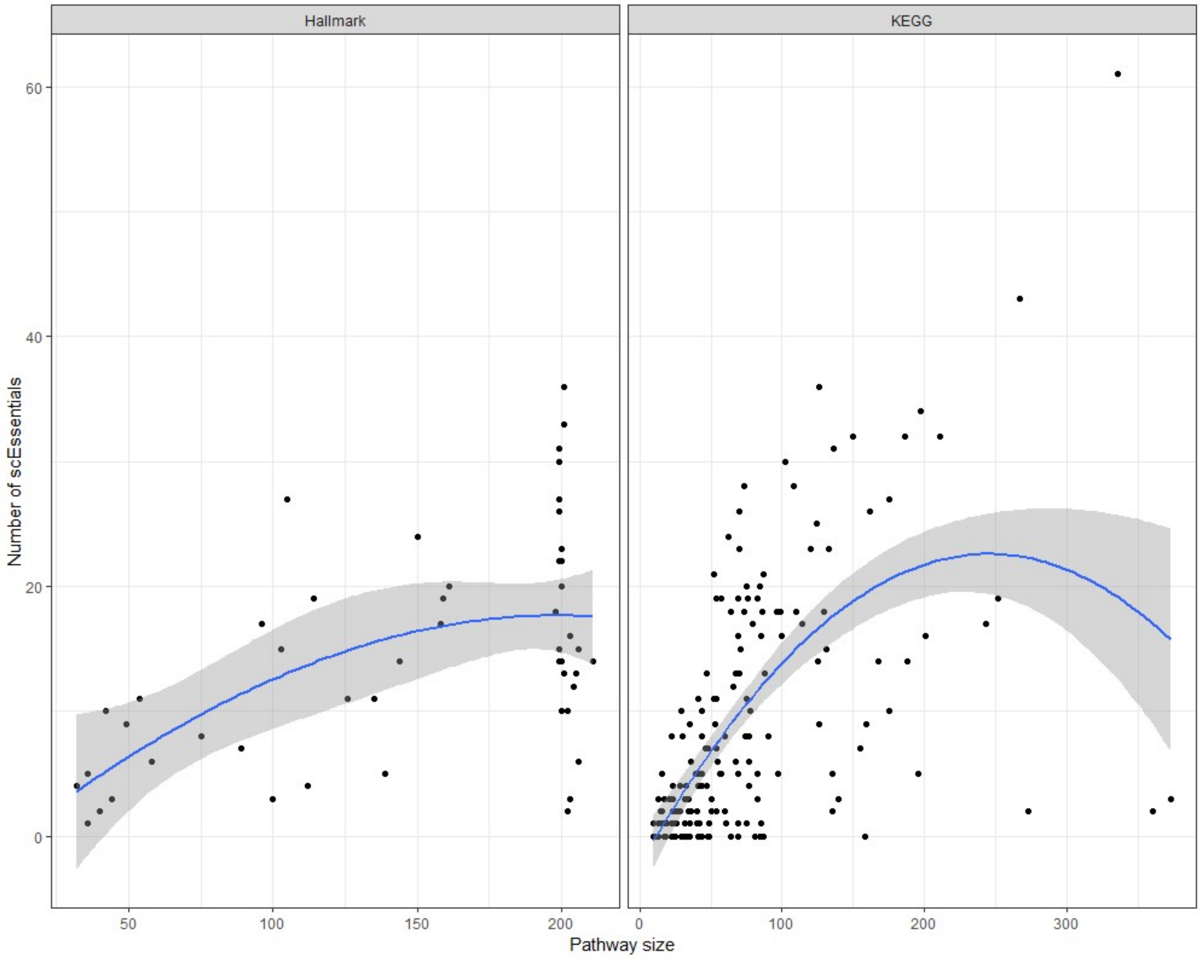


Supplementary Figure 12. The number of scEssentials genes overlapped with each pathway in the Hallmark database (left) and KEGG pathways (right). Linear regression was applied and the standard errors were shaded in grey.

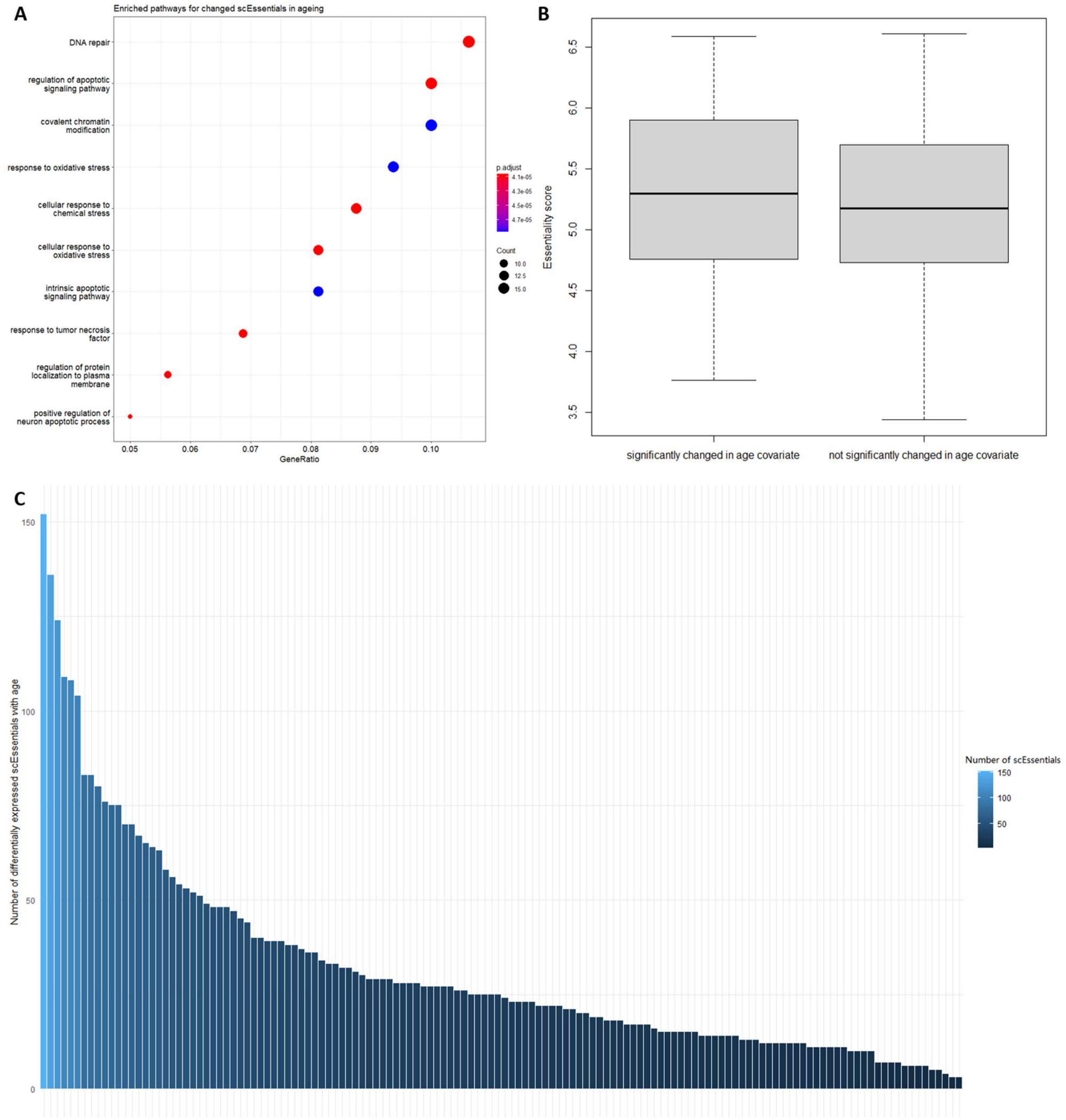


Supplementary Figure 13. Dysregulation of scEssentials genes among different cell types in ageing. A) Dotplot for the top 10 significantly enriched pathways for the scEssentials that changed during age covariates. The analysis was performed with GO database biological processes. B) Boxplot illustrated the significant difference in the essentiality score between significant age covariates and the non-significant age covariate group. Wilcoxon ranked test was applied to determine the significance. C) Barplot showed the number of significantly changed scEssentials across cell types in ageing.
